# Redox control of antibiotic biosynthesis

**DOI:** 10.1101/2025.03.04.641395

**Authors:** Rebecca Devine, Katie Noble, Clare Stevenson, Carlo de Oliveira Martins, Gerhard Saalbach, Hannah P. McDonald, Edward S. Hems, Barrie Wilkinson, Matthew I. Hutchings

**Author notes:** These authors contributed equally to this work (Rebecca Devine, Katie Noble).

## Abstract

*Streptomyces* bacteria make diverse specialised metabolites that form the basis of ∼55% of clinically used antibiotics. Despite this, only 3% of their encoded specialised metabolites have been matched to molecules and understanding how their biosynthesis is controlled is essential to fully exploit their potential. Here we use *Streptomyces formicae* and the formicamycin biosynthetic pathway as a model to understand the complex regulation of specialised metabolism. We analysed all three pathway-specific regulators and found that biosynthesis is subject to negative feedback and redox control via two MarR-family proteins while activation of the pathway is dependent on a cytoplasmic two-component system. Like many *Streptomyces* antibiotics, formicamycins are only produced in solid culture and biosynthesis is switched off in aerated liquid cultures. Here, we demonstrate that a redox-sensitive repressor named ForJ senses oxygen via a single cysteine residue that is required to repress formicamycin biosynthesis in liquid cultures.

## Introduction

New antibiotics are urgently needed to fight emerging drug-resistant infections and specialised metabolites such as those from *Streptomyces* species present a promising source of new molecules ^1^. However, only 3% of all biosynthetic pathways encoded by *Streptomyces* species have been matched to molecules, presumably in part due to a lack of activating signals to induce their expression ^2^. In this work, we aimed to further understand antibiotic biosynthesis by characterising the three cluster-situated regulators in the formicamycin biosynthetic gene cluster (BGC) in *Streptomyces formicae*. Namely we wanted to understand how formicamycin biosynthesis is activated and repressed in response to physiological conditions. Formicamycins represent a new structural class of antibiotics that are potent inhibitors of Gram-positive bacteria including methicillin-resistant *Staphylococcus aureus* (MRSA) and vancomycin-resistant enterococci (VRE), and they have a high barrier for the selection of resistant isolates. The formicamycin BGC (*for* BGC) encodes two MarR-family regulators, ForJ and ForZ, and a two-component system called ForGF in which the sensor kinase (SK) ForG is predicted to be cytoplasmic and activate its cognate response regulator (RR) ForF via phosphorylation.

MarR-family regulators are well known to be involved in the regulation of antibiotic biosynthesis in *Streptomyces* species, e.g., daptomycin production in *Streptomyces roseosporus* is regulated by DptR3 while SAV4189 regulates avermectin production in *S. avermitilis* ^3,4^. They are homodimeric proteins where each monomer contributes a winged helix-turn-helix DNA binding motif, and they often repress transcription of their target genes by binding to palindromic DNA sequences within target gene promoters ^5^. MarR regulators often regulate their own expression and the expression of a divergently encoded gene, although regulation of distally located genes can also occur. Their target gene products have many cellular functions, including antibiotic transport, metabolism, stress responses, virulence and specialised metabolism. MarR proteins typically bind small molecule ligands or phenolic compounds in addition to DNA and, upon ligand binding, there is a conformational change in the transcription factor that alters its interaction with DNA and affects target gene expression ^6^.

Two-component systems typically consist of an SK and its cognate response regulator (RR), encoded within a single operon, that is usually autoregulated. The SK is typically transmembrane and perceives a signal outside the cell and then undergoes a conformational change that activates its kinase domain to autophosphorylate a conserved histidine residue. The cognate RR then catalyses the transfer of the phosphate to a conserved aspartate residue on its own receiver domain. RRs are usually DNA binding proteins and phosphorylation activates them to bind to their DNA targets and modulate target gene expression. Most SKs are bifunctional and when the signal dissipates, they dephosphorylate their cognate RRs and switch off the response^7^.

We previously reported that ForJ is the major repressor of formicamycin biosynthesis and used ChIP-Seq to show that it binds to multiple promoters in the *for* BGC to repress gene expression. Deletion of *forJ* increased compound production more than six-fold, induced the production of new formicamycin congeners, and switched on formicamycin production in liquid culture, something that does not occur in the wild-type strain ^8^. In contrast, ForGF is required for activation of formicamycin biosynthesis, with ForF binding to a single site between the divergent *forGF* and *forHI* genes and activating the expression of both operons. It binds to only one other site on the *S. formicae* genome, upstream of the *KY5_0375* gene, which encodes a putative NLP/P60 family protein that is not required for formicamycin biosynthesis ^9^. ForZ was predicted to regulate the expression of the divergently encoded transporter gene, *forAA*, due to the presence of a specific ChIP-Seq peak at this intergenic promoter (ChIP-Seq data accession no. E-MTAB-8006). However, deletion of *forZ* resulted in a decrease in formicamycin biosynthesis and a decrease in expression of the core biosynthetic genes, even in the absence of ForZ binding to these gene promoters ^8^.

In this work, we aimed to further understand the regulation of formicamycin biosynthesis, and the interplay of ForGF, ForJ and ForZ in controlling this pathway. To achieve this, we used a surface plasmon resonance (SPR) technique called Reusable DNA Capture Technology (RedCaT) to define the exact binding sites of the ForF, ForJ and ForZ regulators ^10^. The position of the ForZ binding site between the divergent *forZ-forAA* genes suggests that ForZ represses expression of *forAA* by binding across its transcription start site. We further show that this is dependent on the concentration of formicamycin congeners present inside the cell because formicamycin binds to ForZ and inactivates its DNA binding activity which likely induces the expression of *forAA*. We confirm that ForF has only three binding sites on the *S. formicae* genome, two upstream of *KY5_0375* and a single site between the divergent *forGF-forHI* operons. The signal for the ForG sensor kinase is not known but we provide evidence that this SK is cytoplasmic and required to phosphorylate and activate ForF. Finally, we show that ForJ binds upstream of all the biosynthetic genes, consistent with ForJ repressing formicamycin biosynthesis. Crucially, we also show that ForJ is redox sensitive, via a single cysteine residue in each monomer, and that dimeric ForJ forms multimers in response to oxidation of these cysteine residues. This provides strong repression of formicamycin biosynthesis under oxidizing conditions and explains why formicamycin biosynthesis is switched off when *S. formicae* is grown in shaking liquid cultures. Together, these regulators balance levels of formicamycin biosynthesis and export in response to multiple environmental signals.

## Results

### ForGF acts as a classical two-component system but with a cytoplasmic sensor kinase

ForGF looks like a classical two-component system, with predicted phosphorylation sites at D53 on the ForF RR and H175 on the ForG SK, but AlphaFold3 suggests ForG does not contain any transmembrane domains (**Fig. S1**) ^11^. Overproduction and purification of the full length His-tagged ForG protein from *E. coli* is consistent with this protein being cytoplasmic with N- and C-terminally 6xHis-tagged proteins both identified in soluble purification fractions (**Fig. S2**). In previous work it was shown that deletion of *forGF* abolishes formicamycin biosynthesis, suggesting that either ForG, ForF or both are required for activation of the pathway ^8^. To test this further, individual Δ*forG* and Δ*forF* mutants were made in *S. formicae* and complemented by introduction of the wild-type genes *in trans* in single copy under the control of the native *forGF* operon promoter. Measurement of the levels of formicamycins show that loss of either ForG or ForF abolishes their biosynthesis, which is restored by introducing a replacement copy, *in trans* (**Fig. 1**). Next, the codons for H175 in *forG* and D53 in *forF* were changed to the codon for alanine, to make H175A ForG and D53A ForF variants which cannot be phosphorylated. The D53 in ForF was additionally changed to glutamate to make a D53E variant of ForF as such substitutions have been shown to mimic the phosphorylated aspartate for some RRs ^12^. The alterations were achieved through both single point mutations of the native alleles using CRISPR/Cas9, and by complementing the single gene deletion mutants *in trans* but with the single point mutants instead of the native genes. Strains producing the phosphodeficient variants of ForG or ForF showed a near complete loss of formicamycin biosynthesis while the strain producing the phosphomimetic D53E variant of ForF showed production levels that were higher than that of the wild-type strain (**Fig. 1**). These data support the hypothesis that ForGF acts as a typical two-component system, i.e., ForG phosphorylates and activates ForF, and that D53E ForF is phosphomimetic.

**Figure 1.**
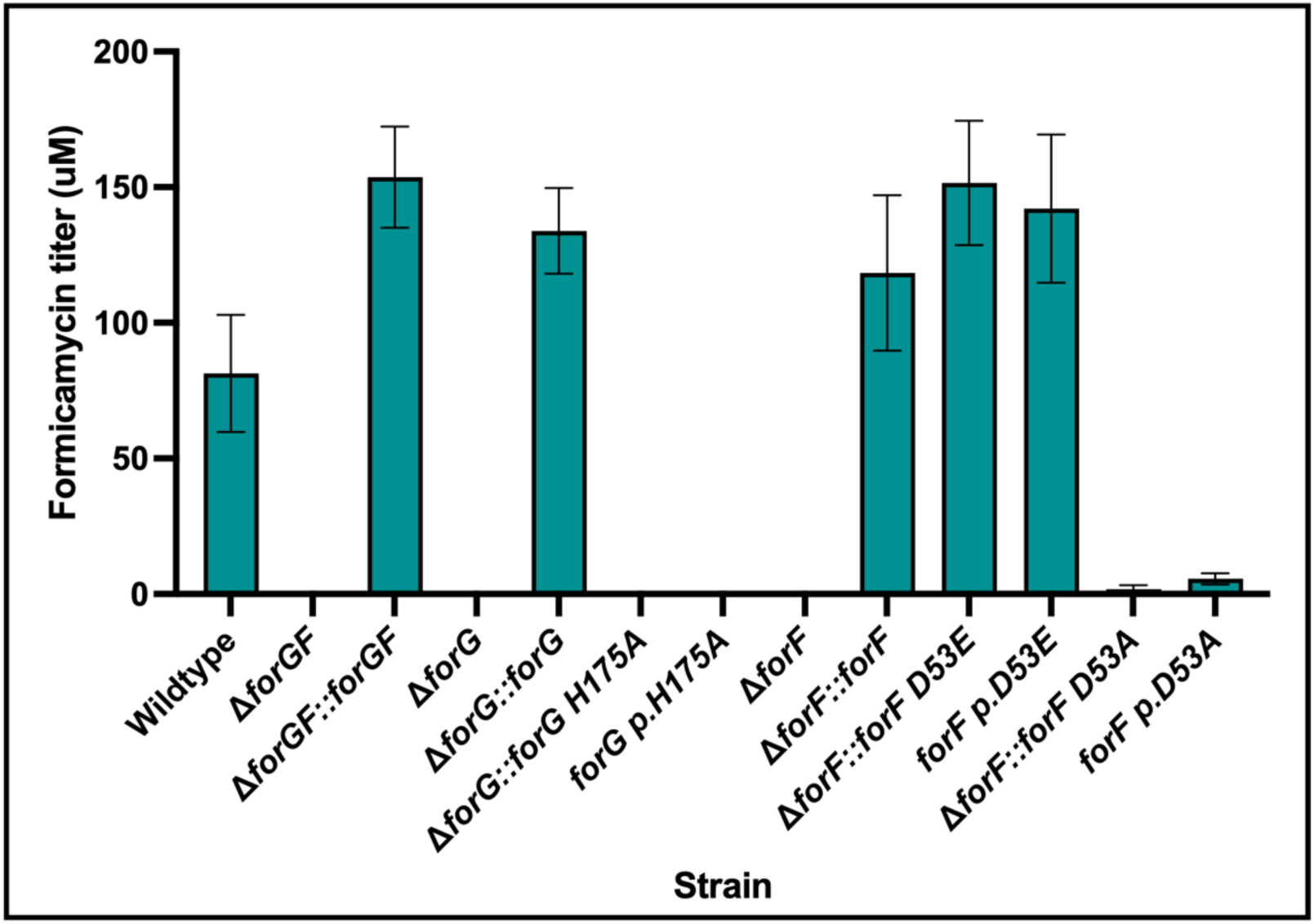
ForGF acts as a classical two-component system to activate formicamycin biosynthesis. Deletion of either *forF* or *forG* individually, or the entire *forGF* operon, abolishes formicamycin production, which can be restored by complementation *in trans* under the native *forGF* operon promoter. Changing the conserved histidine residue of ForG (H175) or the conserved aspartate residue of ForF (D53) to an alanine via either *in trans* complementation or introduction of point mutations directly into the native chromosomal alleles by gene editing (p.) results in a near total loss of biosynthesis. Changing the D53 of ForF to a glutamate residue by gene editing (p.) appears to mimic the phosphorylated D53 residue and restores formicamycin biosynthesis.

To investigate its DNA binding properties *in vitro*, ForF was over-produced and purified from *E. coli* for Reusable DNA Capture Technique (ReDCaT) SPR experiments, in which overlapping tiled oligonucleotide probes are designed against a target promoter to identify the exact transcription factor binding site ^10^. ChIP-Seq analysis revealed previously that ForF binds to a single site in the formicamycin BGC, between the divergent *forGF* and *forHI* genes, and reporter assays indicated that the *forGF* and *forHI* promoters are both activated by ForF ^8^. These ChIP-Seq data also showed a second binding site of ForF upstream of the *KY5_0375* gene, which encodes a putative NLP/P60 family protein ^8^. To determine the ForF binding site relative to the transcription start sites of these operons and gene, we designed double stranded oligonucleotide probes to span this entire region and found that ForF binds to a single site between the divergent *forHI-forGF* operons and to two sites in the region upstream of *KY5_0375*. Further truncation of each of these probes identified a 12bp consensus sequence that is required for ForF binding (**Fig 2**).

**Figure 2.**
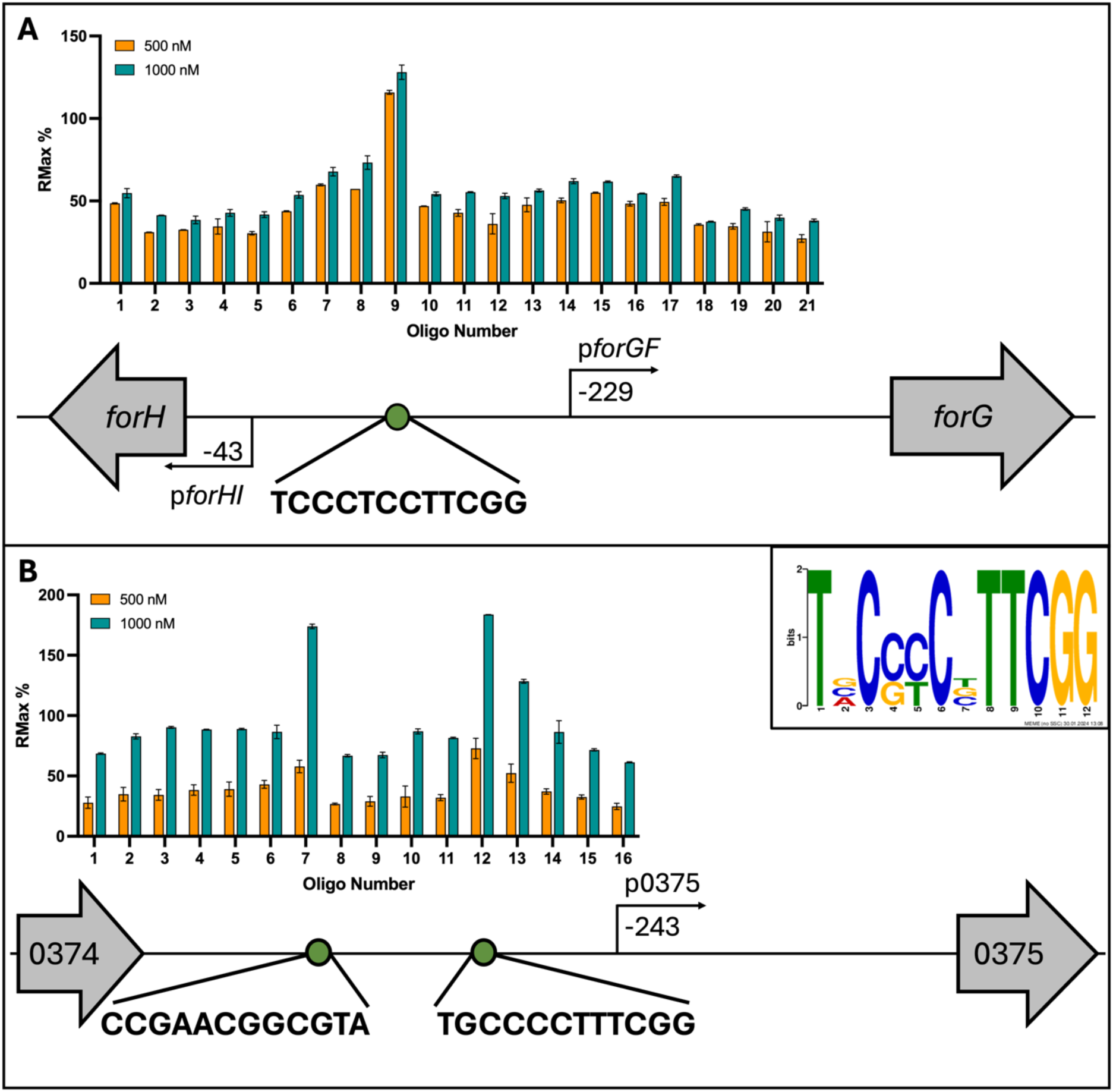
ForF binds to a 12 base pair DNA sequence at just three sites on the *S. formicae* genome. **A.** ForF binds to a single site between the divergent *forHI* and *forGF* operons. **B.** ForF binds to two sites upstream of the *KY5_0375* gene. **Inset.** A consensus sequence generated by aligning these three binding sites. Distances between the start of the coding regions, promoters and binding sites are scaled.

### ForZ blocks the transcription start site of the exporter gene, *forAA,* and autoactivates its own expression

Previously, ChIP-Seq experiments showed that ForZ binds to a single region in the *S. formicae* genome, between *forZ* and the divergently encoded gene *forAA* gene, which likely encodes a transporter for fasamycins and formicamycins ^8^. To confirm ForZ binding to this region the ForZ protein was over-produced and purified from *E. coli* and used in ReDCaT SPR reactions to identify interactions with double stranded, overlapping DNA probes that span across the intergenic region between *forZ* and *forAA*. The results show that ForZ specifically binds to a single DNA probe and using sequential truncation, we identified the ForZ DNA binding site within this probe as an imperfect palindrome with an 8bp repeat sequence, **TAGCTCG<u>A</u>AGTTCGAT**, where **<u>A</u>** is the transcription start site (TSS) of the *forAA* gene. Consistent with the ChIP-seq data, this DNA sequence is not present anywhere else in the *S. formicae* genome, confirming there is a single ForZ binding site in *S. formicae*, within the *forZ-forAA* divergent promoters (**Fig 3**). The location of this binding site is such that binding of ForZ would repress the expression of *forAA* by occluding the TSS to block RNA-polymerase binding. However, expression of the divergent *forZ* gene is leaderless, such that ForZ binding would overlap its -35 region and thus ForZ could autoactivate its own expression.

**Figure 3.**
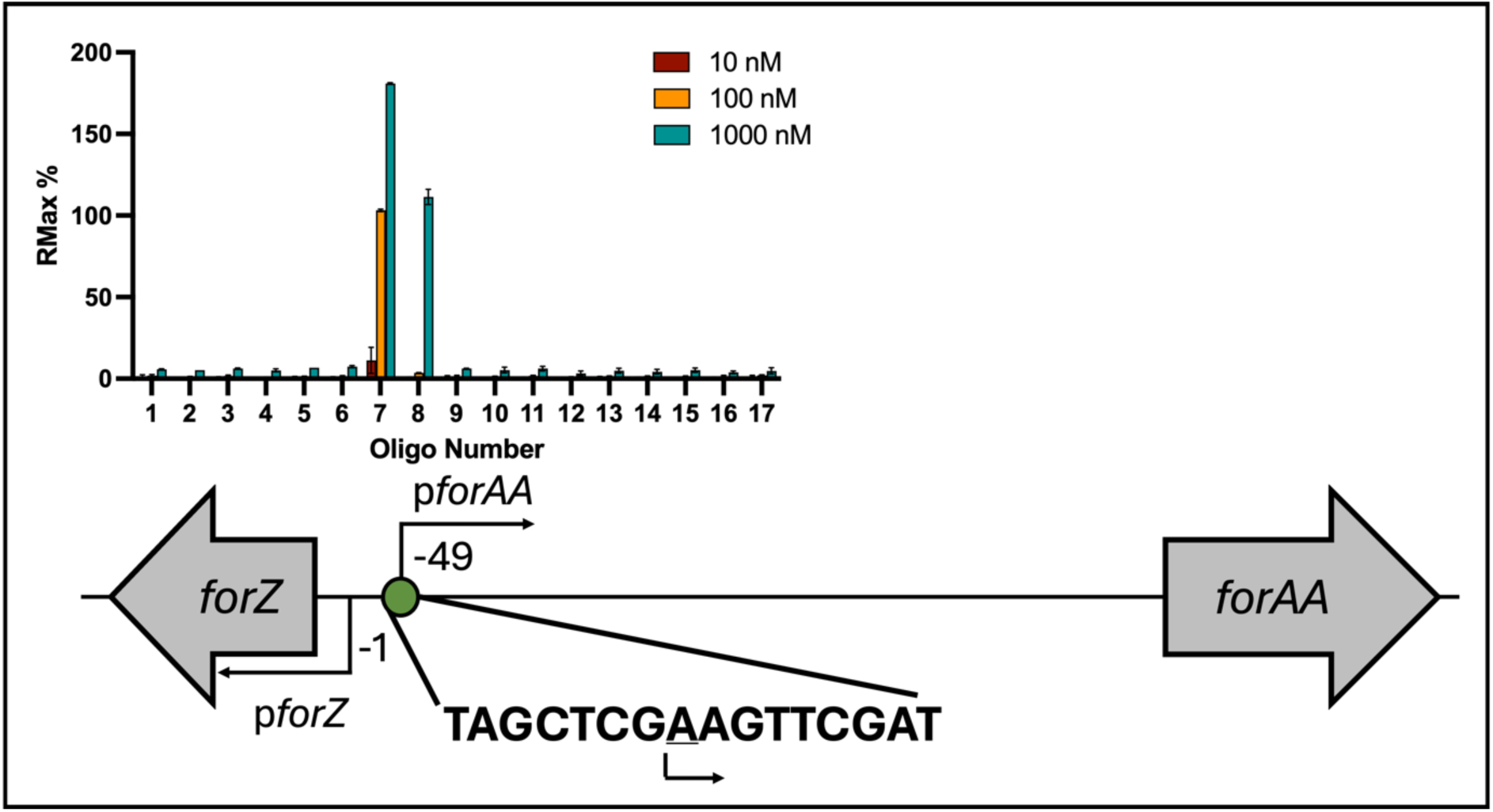
ForZ binds an imperfect inverted repeat to repress expression of *forAA.* ForZ specifically binds to an 8 base pair palindromic repeat sequence between the divergent *forZ* and *forAA* genes in a concentration-dependent manner, where the underlined A is the TSS of *forAA*. Distances between the start of the coding regions, promoters and binding sites are scaled.

### High level formicamycin export leads to pathway shutdown

Our previous work suggested binding of ForZ to DNA might activate *forAA* expression, as levels of the *forAA* transcript are 3-fold downregulated in a Δ*forZ* deletion mutant. However, the SPR results here show that binding of ForZ to the *forAA* promoter must block RNA polymerase access and repress expression of *forAA* because the ForZ binding site overlaps its TSS (**Fig. 3**). Although contradictory, we suggest that in the absence of a controllable export mechanism, the entire formicamycin BGC is downregulated by an alternative feedback mechanism, to prevent metabolically expensive compound overproduction and build up that could be toxic to the cell. In agreement with this, we showed previously that several other *for* BGC transcripts are down-regulated in a *forZ* deletion mutant, and formicamycin titres are reduced to approximately 60% of the wild-type levels in the Δ*forZ* strain, even though ForZ only binds to a single site within the divergent *forZ-forAA* promoter regions ^8^.

### Formicamycin binding to the ForZ protein abolishes its DNA binding activity

MarR-family regulators are commonly involved in regulating genes that control the export of antibiotics, i.e., drug efflux pump genes such as *forAA* ^4,13^. In these instances, the associated ligand that releases the MarR regulator from DNA is often the transporter substrate, i.e., the end product of the pathway. To test whether ForZ binding to the *forAA* promoter is abolished by fasamycins or formicamycins, purified ForZ binding to a DNA probe containing the ForZ binding site (**Fig. 4**) was tested in a concentration-dependent manner in the presence of either 10 mM fasamycin E or formicamycin I (**Fig. 4**) using a multi-cycle affinity kinetics protocol. The addition of fasamycin E to the binding reactions had a small effect on DNA binding, except at the highest protein concentrations, where ForZ binding to DNA was reduced to around 50% of the normal R_max_. However, the addition of formicamycin I completely abolished binding of ForZ to the DNA (**Fig. 4**). This suggests formicamycins, and to a lesser extent fasamycins, are the ligands that bind to ForZ and cause a conformational change that reduces its ability to bind DNA thus inducing *forAA* expression. By titrating increasing concentrations of formicamycin I into binding reactions between ForZ (500 nM) and DNA (1 mM), we identified the IC_50_ of this interaction to be 2.48 μM (**Fig. 4**).

**Figure 4.**
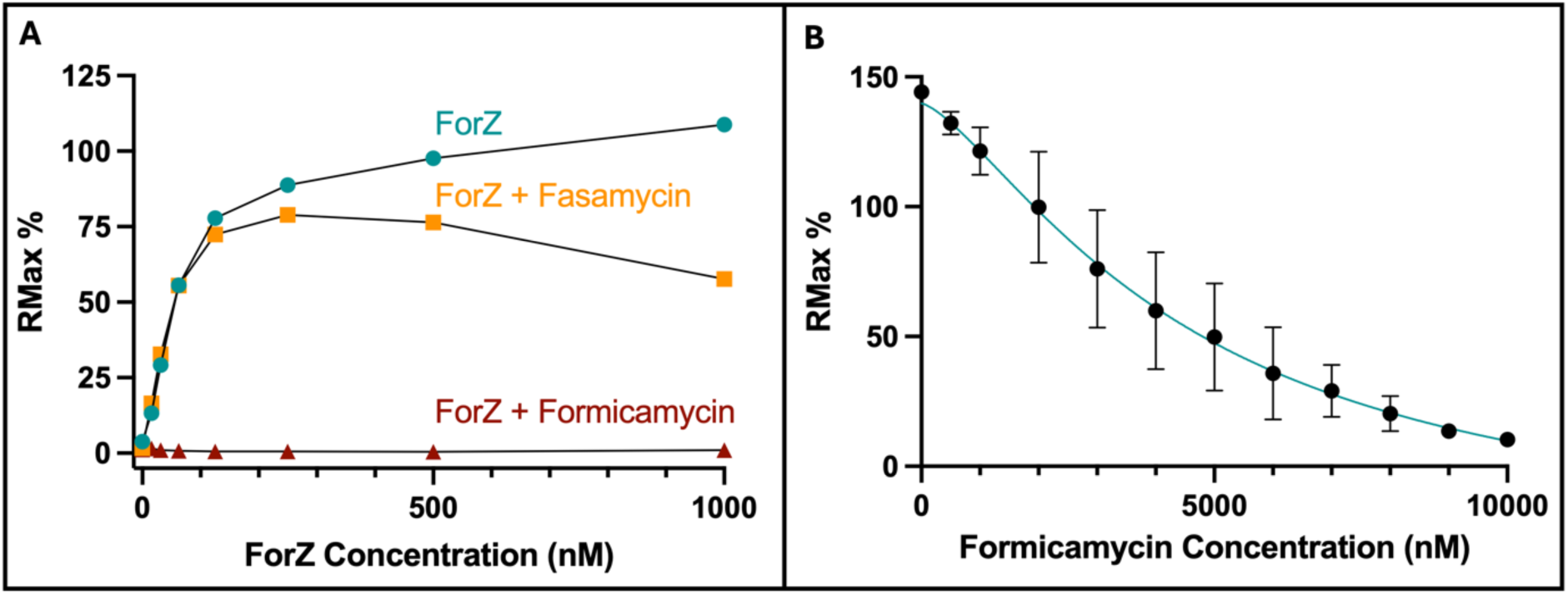
ForZ binding to DNA is abolished by the addition of formicamycin I. **A.** SPR was used to measure the interaction of a double stranded DNA probe containing the inverted repeat sequence (AT<u>TAGCTCGAAGTTCGAT</u>GCATCTTGCAGT) with increasing concentrations of ForZ protein. ForZ binds to DNA with a KD of 46 nM (**Fig S3**.). Addition of formicamycin I abolished the ability of ForZ to bind to DNA, and fasamycin E reduces binding to approximately 50% at 10 µM (n=2) **B.** Titrating increasing concentrations of formicamycin I onto 500 nM ForZ shows that formicamycin I inhibits ForZ binding to DNA with an IC50 of 2.48 μM (n=4).

These data support a hypothesis in which ForZ exists preferentially in its DNA bound form, because the K_D_ of the interaction between ForZ and DNA is considerably lower than the IC_50_ at which formicamycin I inhibits this interaction. It also supports the hypothesis that ForZ regulates the export of formicamycins after they are synthesized and begin to accumulate inside the cell. In *S. formicae,* this would ensure that *forAA* expression is repressed until formicamycin concentrations reach a certain threshold.

### ForJ represses the formicamycin biosynthetic genes and forms multimers in oxidizing conditions

Our previous work has indicated that ForJ binds at multiple locations across the formicamycin BGC where it represses the expression of the biosynthetic transcripts. To confirm ForJ binding at these promoters, double stranded oligonucleotide probes were designed against the intergenic region between the divergent *forT* and *forU* genes and used to measure ForJ binding using ReDCaT SPR. Sequence specific binding of ForJ was only observed in the presence of a reducing agent (1 mM DTT), suggesting ForJ is redox-sensitive. Probes were then designed against other regions within the *for* BGC that were enriched in the ForJ ChIP-seq experiment, namely the *forM, forGF* and *forJ* promoters and the end of the coding region of *forE*. Despite the presence of ChIP peaks, no ForJ binding was observed at the *forGF* promoter or in the coding region of *forE* in these experiments. However, sequence specific binding was observed at the *forM* and *forJ* promoters and alignment of the sequences bound at each promoter (divergent *forT*-*forU*, divergent *forN-forM* and *forJ*) was used to identify the consensus binding site of ForJ, which is a 14 base pair A/T rich sequence that spans the TSS of the *forM*, *forJ* and *forT* genes. The *forM* promoter contains two ForJ binding sites, back-to-back, suggesting that two ForJ dimers could bind this region (**Fig. 5**). Multi-cycle affinity kinetics allows several concentrations of ForJ to be assessed for binding affinity to the DNA sequence, with chip regeneration between each cycle. The results show that ForJ binds this consensus sequence relatively weakly, at least in the presence of DTT, with a K_D_ of 288 nM (**Fig. S3**).

**Figure 5:**
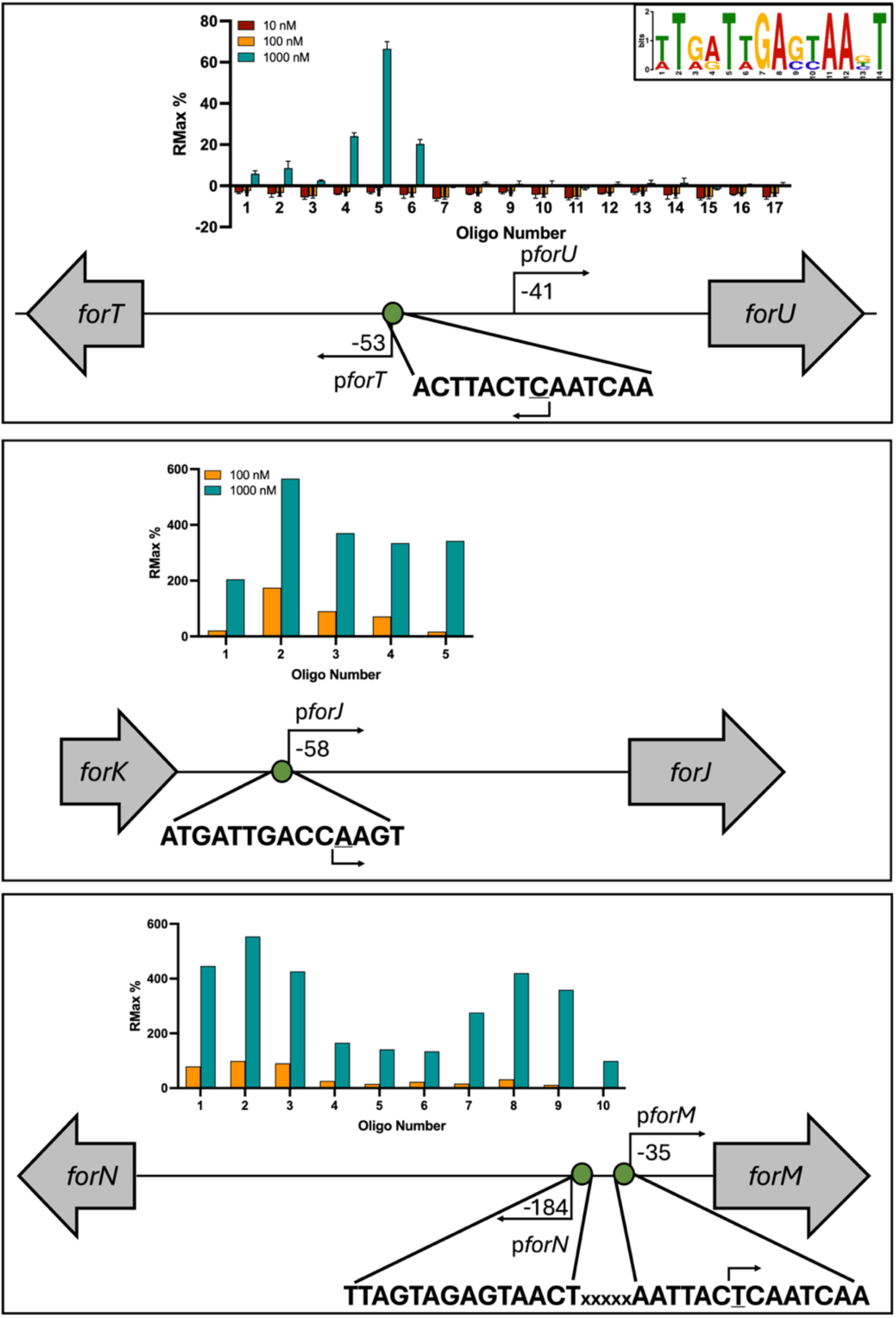
ForJ binds upstream of several biosynthetic genes in the *for* BGC. ForJ binds specific sequences within the promoters of *forM*, *forJ* and *forT* in the presence of 1 mM DTT, and alignment of the binding sites identified a 14 bp consensus site (inset). The site appears twice, back-to-back in the promoter of *forM*. Binding at the *forT/*U promoter was tested three times at three concentrations, whereas binding at the other sites was tested twice. Previous work shows binding of ForJ to these promoter regions inhibits gene expression (Devine *et al.*, 2021) likely because ForJ blocks access to their TSS (underlined). Distances between the start of coding region, promoters and binding sites are scaled.

There are several published examples of redox sensitive MarR-family regulators, including HypR in *Bacillus subtilis,* CosR in *Corynebacterium glutamicum* and AbfR in *Staphylococcus epidermidis* ^14–16^. Typically, redox sensitivity is mediated through a surface exposed cysteine residue that can become oxidized and change the conformation of the MarR protein to affect its DNA binding activity. ForJ contains a single cysteine residue at position 68 and, according to AlphaFold models, this residue is in the DNA binding domain of the protein (**Fig. 6**). To investigate what type of structural change might be occurring upon oxidation of C68, ReFeyn mass photometry was used to measure the size of both reduced and oxidized ForJ. The mass of reduced ForJ is around 37 kDa, the expected size of a ForJ dimer, whereas upon oxidation, the mass shifts to around 75 kDa. This was also observed on SDS-page gels when ForJ was exposed to increasing concentrations of either a reducing agent (DTT) or an oxidizing agent (H_2_O_2_) and analysed using SDS-PAGE, omitting the addition of β-mercaptoethanol and boiling of samples prior to loading to prevent the reduction of disulphide bonds (**Fig. 6**). These results are consistent with two ForJ dimers binding to each other upon oxidation, by forming inter-subunit disulphide bonds between the cysteine residues on adjacent dimers, as has been reported for other redox-sensitive MarR-family regulators ^17,18^. Indeed, SPR reactions with oxidized ForJ showed extremely strong binding to all tested DNA probes, which may suggest that multimers were forming during the reactions. Sequence specific binding was only observed after the addition of DTT. AlphaFold3 modelling suggests the cysteine residue is not required for protein folding and in agreement with this, we were able to express a 3xFLAG-tagged ForJ C68S protein in *S. formicae* and confirm its presence by western blot analysis, suggesting the cysteine residue is not essential for protein folding in the native host (**Fig. S4**). However, we were unable to over-produce and purify the C68S protein from *E. coli*.

**Figure 6:**
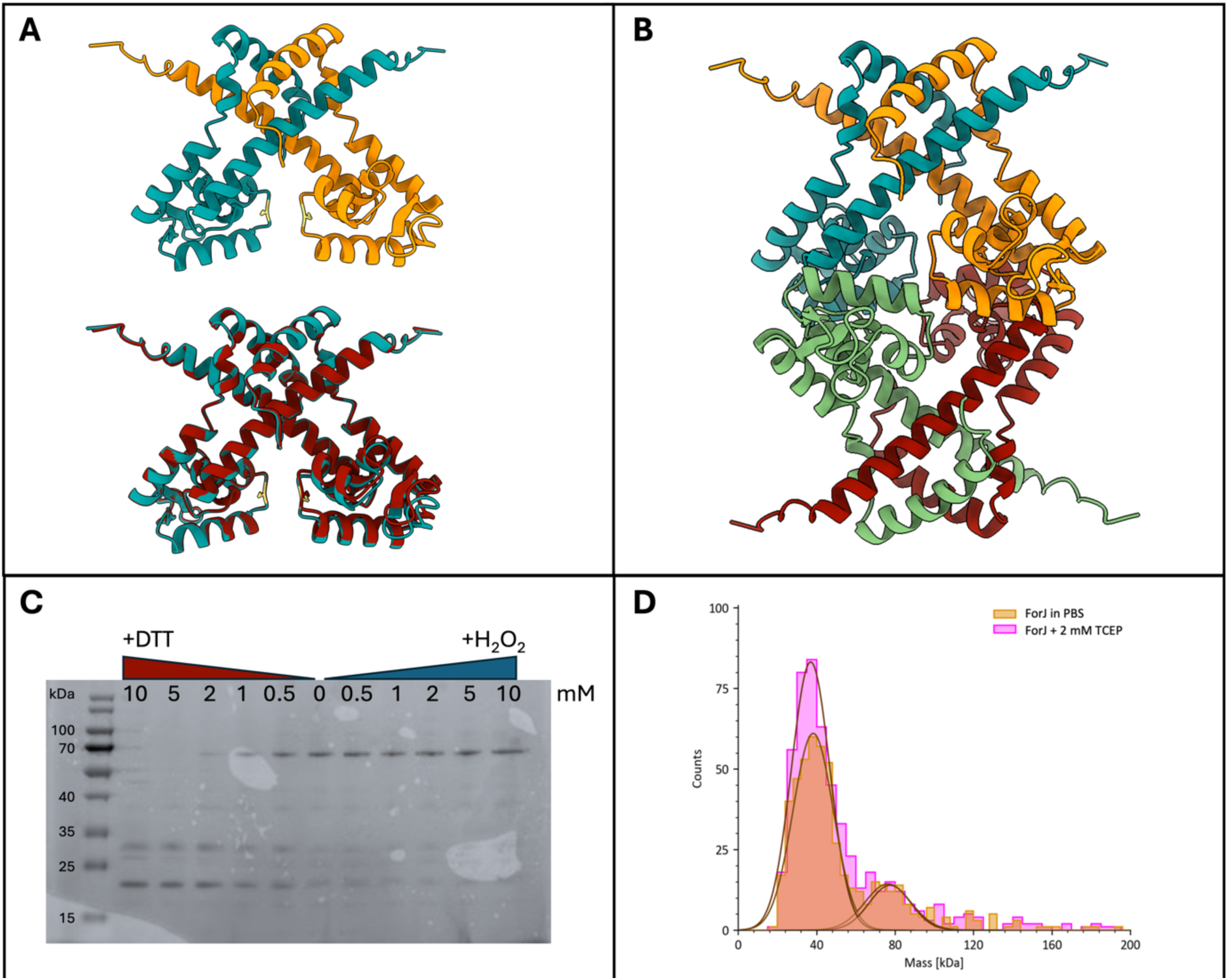
ForJ forms dimers-of-dimers in oxidizing conditions *in vitro.* **A:** Top: AlphaFold 3 model of a wild-type ForJ dimer, with the monomers in blue and orange. Bottom: AlphaFold model of the the C68S variant dimer (red) overlaid with the wild-type (blue) suggesting the cysteine residue (yellow) is not essential for protein folding. Note that C68 is predicted to be in the DNA binding domain of ForJ but in dimeric ForJ the cysteines are facing in opposite directions. **B:** AlphaFold 3 model of wildtype ForJ dimer of dimers. This suggests that on oxidation, two dimers of ForJ could tightly wrap around the DNA, providing strong repression of gene expression. **C:** Analysis by non-denaturing SDS-PAGE shows the size of ForJ increases upon exposure to hydrogen peroxide consistent with a dimer of dimers (76 kDa) formed by oxidation of the cysteine residues, but it forms single dimers (38 kDa) and monomers (19 kDa) in the presence of the reducing agent DTT. **D:** ReFeyn mass photometry shows that in the absence of reducing agent, native ForJ is present in two states; 38 kDa and 76 kDa. Reduction by TCEP keeps the majority of the sample at 38 kDa, the expected size of a ForJ dimer. Mass photometry experiments were run by Colin Grant, ReFeyn Ltd., Oxford.

### *S. formicae* down-regulates formicamycin production under oxidizing conditions

To determine whether oxidation of ForJ C68 affects DNA binding *in vivo,* we used qRT-PCR to measure the expression of the *for* BGC in *S. formicae* grown on agar in the absence and presence of 0.5 mM diamide, which specifically catalyses disulphide-bond formation in thiol-containing proteins. The results show that, with the exception of *forGF*, the levels of gene expression across the *for* BCG are approximately halved in the presence of diamide compared to *S. formicae* grown in standard growth medium (**Fig. 7**). These results suggest the *for* BGC is repressed under oxidizing conditions. We propose this is due to ForJ binding more tightly to DNA by forming dimers-of-dimers when C68 is oxidized, as seen *in vitro*. Indeed, this could explain why we see ChIP-seq peaks throughout the *for* BGC even in the absence of true ForJ binding sites, i.e., ForJ is winding up the DNA. This would further block transcription resulting in a shut down in formicamycin biosynthesis.

**Figure 7:**
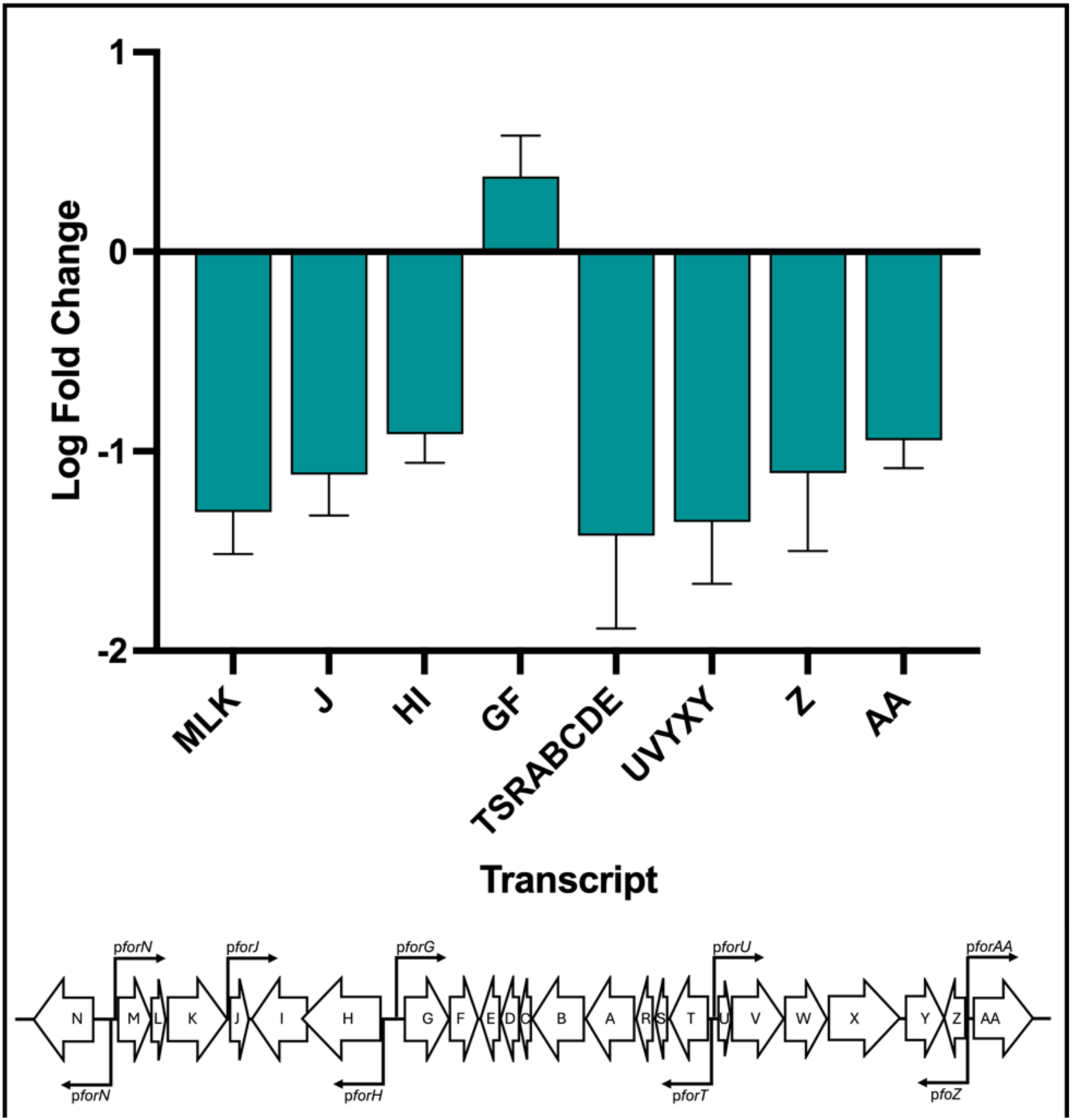
Growing *S. formicae* under oxidizing conditions results in down-regulation of the *for* BGC and a decrease in formicamycin production. Addition of 0.5 mM diamide to the solid growth medium down-regulates expression of *for* BGC transcripts encoding the biosynthetic and export machinery (n=4).

In previous work, we noted that wild-type *S. formicae* only produces formicamycins on solid media, and deletion of *forJ* is required to induce production in liquid cultures ^8,9^. Considering the data presented here, we propose that de-repression of the *for* BGC in liquid cultures of *S. formicae* Δ*forJ* is due to the loss of redox sensitivity of ForJ, as liquid cultures are more aerobic environments than solid cultures, with more headspace and shaking to increase aeration. To investigate this, we mutated *forJ* in wild-type *S. formicae*, to encode a ForJ C68S variant, and tested the expression levels of *for* BGC transcripts as well as the ability of this mutant to produce formicamycin congeners in liquid culture. *S. formicae* carrying the ForJ C68S allele was found to over-express all *for* BGC transcripts to a similar extent to the *ΔforJ* mutant, and both strains show a significant up-regulation compared to the wild-type strain in liquid culture. The only transcript that is down-regulated in both mutants is the *forZ* transcript, but this is consistent with increased expression of *forAA* to cope with the increase in compound production which would also block autoactivation of *forZ* expression. Strikingly, like *S. formicae* Δ*forJ, the S. formicae* ForJ C68S mutant can produce fasamycins and formicamycins in liquid culture, indicating that this single cysteine residue is responsible for repression of the pathway under oxygen rich conditions (**Fig. 8**). These data support the hypothesis that, in wild-type *S. formicae*, ForJ is predominantly in the dimer-of-dimers form in liquid culture due to increased oxidation of the cysteine residue resulting in stronger repression of the pathway. By deleting *forJ* or making ForJ ‘blind’ to oxidation via substitution of C68, ForJ remains in its dimeric form, meaning binding to the DNA is weaker and there is less repression of *for* BGC expression.

**Figure 8:**
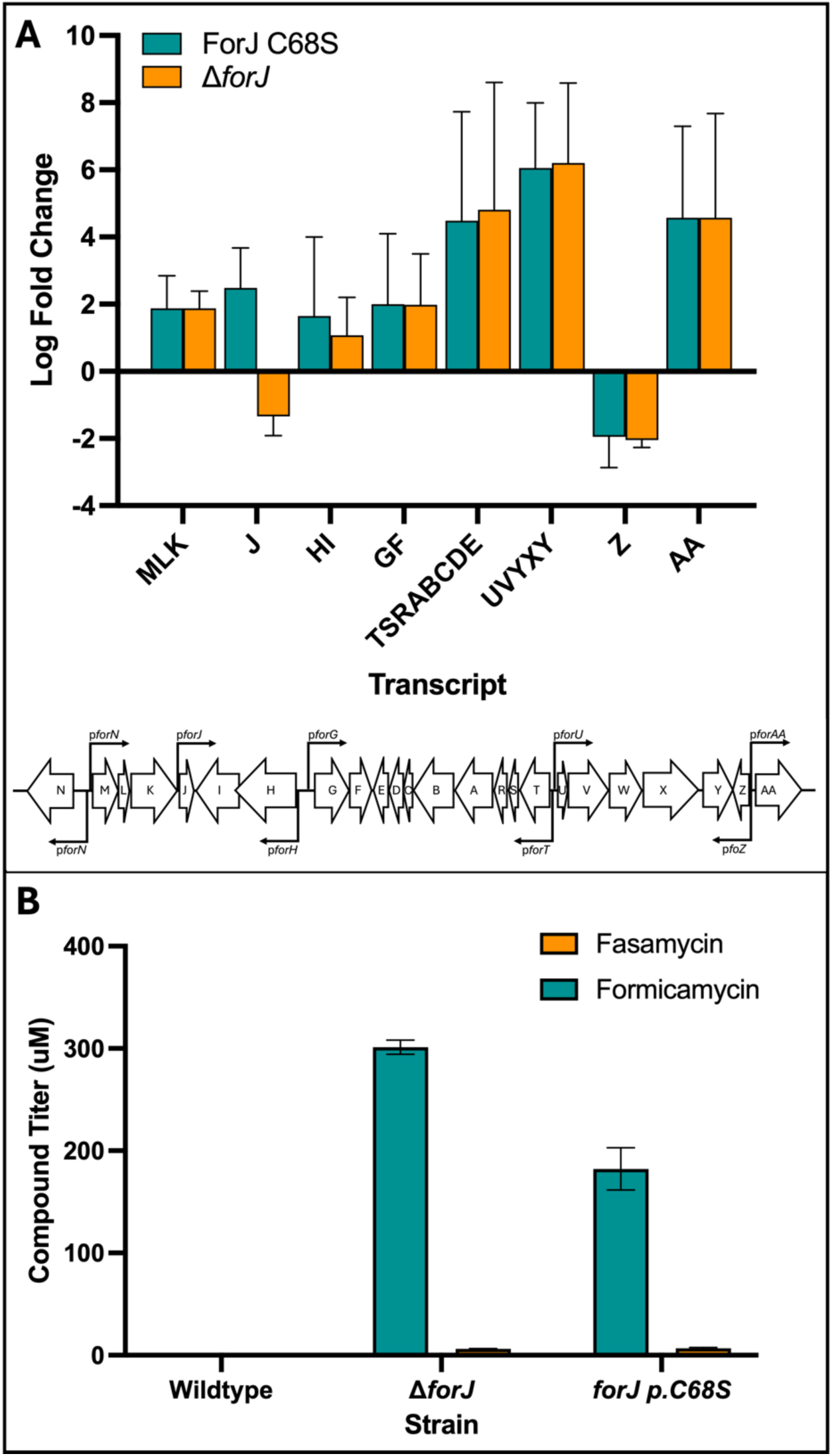
The oxidation state of ForJ C68 controls the expression levels of the *for* BGC. **A:** In liquid culture, qRT-PCR data show that, with the exception of *forZ*, expression of the *for* BGC is increased in both a *forJ* deletion mutant and the ForJ C68S mutant compared to the wild-type (n=3). **B:** Fasamycin and formicamycin congeners are only produced in liquid cultures of *S. formicae* Δ*forJ* and *S. formicae* ForJ C68S, where the native *forJ* allele has been mutated (n=4), and not in the wild-type strain. This suggests the oxidation state of C68 is important for determining whether the pathway is on or off during liquid culture.

### Deletion of *forJ* results in increased expression of proteins involved in formicamycin biosynthesis and stress responses

Some secondary metabolites are redox active due to the presence of quinone groups that can be reduced to the semi-quinone form, e.g., actinorhodin produced by *S. coelicolor* ^19^. Fasamycins and/or formicamycins do not contain the typical features of redox-active metabolites, so the biological purpose of the redox-dependent regulation of the *for* BGC is unclear. However, whole cell proteomics analysis of *S. formicae* Δ*forJ* shows significant up-regulation of proteins encoded by the *for* BGC, as well as various proteases, hydrolases and oxidoreductases, and down-regulation of proteins involved in growth and development, such as cell division proteins, RNA polymerase subunits and proteins involved in the biosynthesis of membrane components (**Fig. 9**). This suggests that increased formicamycin biosynthesis results in cellular stress to the host. The process of biosynthesising a fasamycin and/or formicamycin involves multiple flavin dependent enzymes ^20^, including the halogenase ForV which catalyses up to 5 chlorination reactions on a single formicamycin molecule, each via the intermediary of a molecule of the strong oxidizing agent hypochlorous acid ^21^. Thus, high levels of formicamycin biosynthesis may correlate with the formation of oxidative species that can cause damage to other cellular proteins. We propose that the redox sensitivity of ForJ presents an advantage to *S. formicae* by repressing formicamycin biosynthesis when levels of biosynthesis become so high that these reactive by-products build up, ensuring formicamycin biosynthesis does not cause oxidative stress inside the cell.

**Figure 9:**
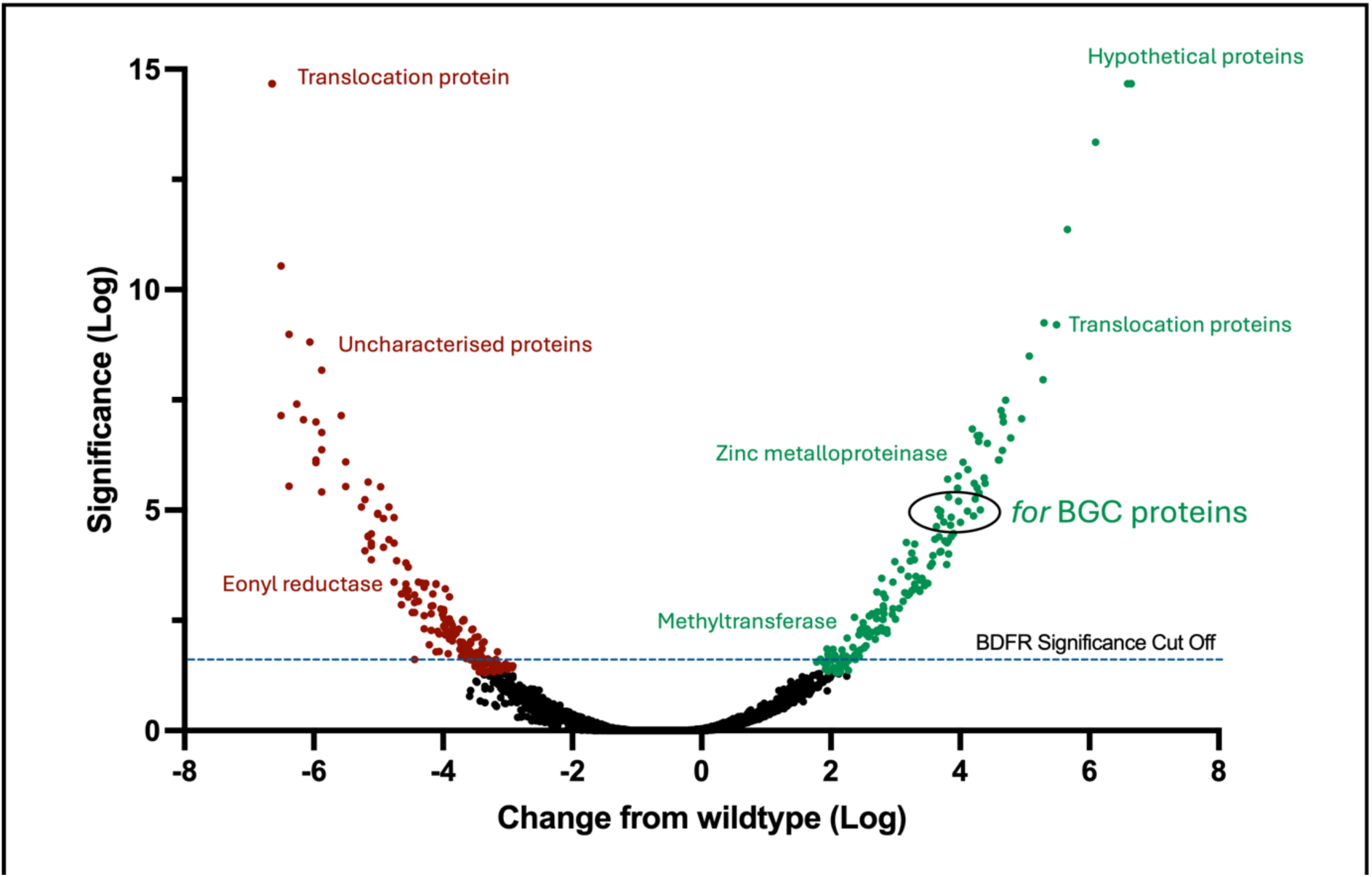
Increased formicamycin biosynthesis is associated with increased production of stress response proteins. Removing the redox-sensitive repressor ForJ (*ΔforJ*) results in significant up-regulation of proteins from the formicamycin biosynthetic pathway as well as stress-response proteins such as those involved in proteolysis. This is consistent with an oxidative-stress response being triggered. In contrast, competing biosynthetic pathways such as PKSs and siderophores, membrane components and cell division proteins are downregulated.

## Discussion

In this work we have characterised all three BGC-situated regulators involved in formicamycin biosynthesis. We have shown that ForGF acts as a classical two-component system to activate biosynthesis in response to an unknown signal(s), but ForG must sense these signals in the cytoplasm because it is cytoplasmic and not membrane bound. We propose that activated ForG phosphorylates ForF which then activates the expression of two genes (*forHI*) involved in boosting levels of the precursor malonyl-CoA, which is required to make the polyketide chain. The other ForF target, *KY5_0375*, cannot be involved in formicamycin biosynthesis because we have shown the pathway only requires the 24 genes in the *for* BGC ^9^.

ForJ is the major repressor of formicamycin biosynthesis and binds a consensus sequence in several key biosynthetic gene promoters. Binding of ForJ to DNA blocks the transcription start sites of these genes, thereby blocking transcription by occluding RNA polymerase binding. Under reducing conditions, e.g., in solid culture, we propose that ForJ is a weak repressor of the *for* BGC, such that biosynthesis occurs at a low level. As formicamycins accumulate, ForZ detects these and releases repression of the exporter gene, *forAA*, which facilitates export of the fasamycins and formicamycins from the cell. This may also confer immunity on the producing organism since there are no other resistance genes within the *for* BGC. This coordination of biosynthesis and export by ForF, ForJ and ForZ ensures maximum conversion of fasamycins to formicamycins before the export mechanism is activated. Under more oxidizing conditions, we propose that inter-subunit disulphides form between cysteine residues on adjacent ForJ dimers, leading to dimer-of-dimers formation, which results in stronger repression of the pathway. We further propose that accumulating biproducts from the conversion of a fasamycin to a formicamycin may be one such cause of this increased oxidative stress, meaning that when levels of biosynthesis get too high, ForJ undergoes structural changes that result in stronger repression of the pathway. In this way, biosynthesis and export are maintained at the optimal levels to ensure they don’t result in toxic accumulation of formicamycins inside the cell (**Fig. 10**).

**Fig. 10.**
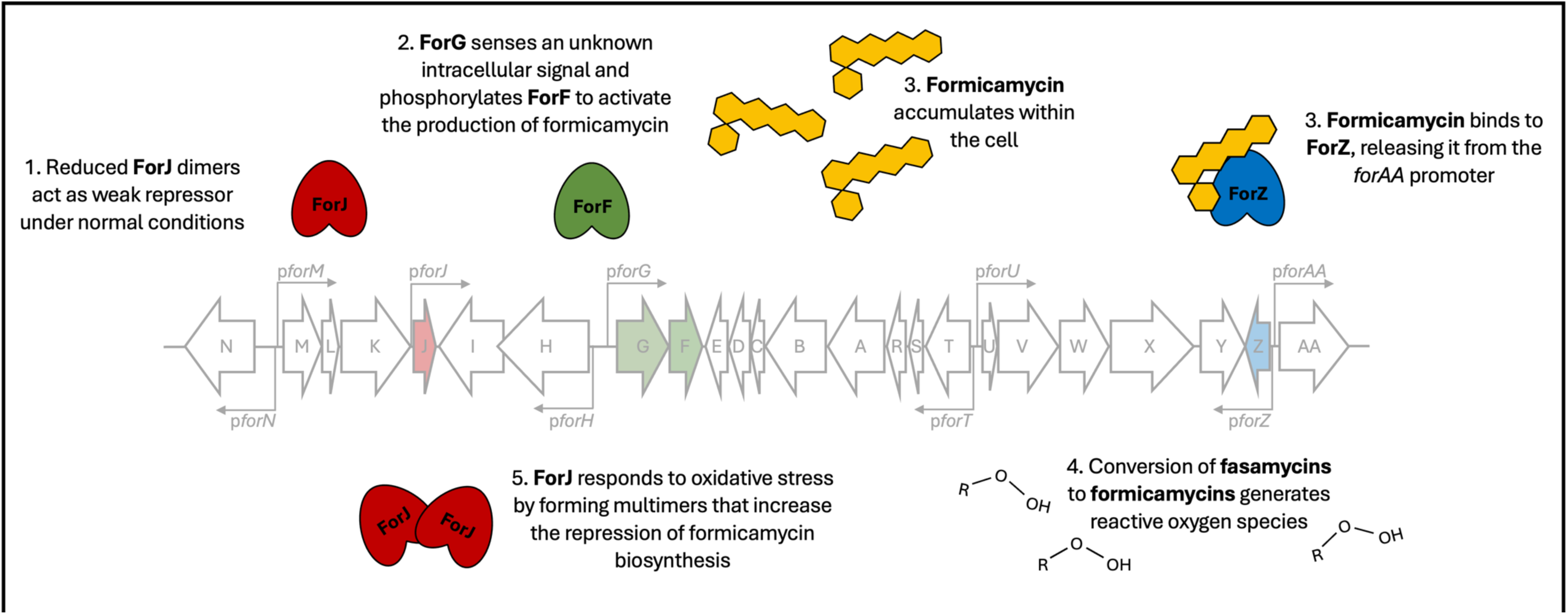
An overview of cluster-situated regulatory control of formicamycin biosynthesis by ForJ (repressor), ForGF (activator) and ForZ (export): Under reducing conditions, ForJ is bound as a dimer to multiple biosynthetic promoters, providing weak repression of the pathway. ForZ is bound to the divergent *pforZ-AA* promoters, repressing expression of the export pump. The kinase ForG senses an unknown intracellular signal and phosphorylates the response regulator, ForF. Once phosphorylated, ForF binds a consensus sequence in the divergent *pforHI-GF* promoters to activate formicamycin biosynthesis. Once formicamycins begin to accumulate, ForZ repression of *forAA* is lifted, allowing for the compounds to be exported from the cell. Increased formicamycin biosynthesis results in the accumulation of oxidative species. Oxidation of a cysteine residue in ForJ results in inter-dimer disulphide bonds forming, resulting in the formation of a dimer of dimers that represses more strongly. Together, these regulators ensure formicamycin biosynthesis and export occur at an optimal rate.

Repression of *forAA* by ForZ is typical of MarR-family regulators, and many characterised examples exist in the literature where MarR proteins repress expression of an export pump until DNA-binding is attenuated by the export substrate ^22^. For example, CtcS, a MarR-family regulator encoded in the chlortetracycline biosynthetic pathway in *Streptomyces aureofaciens,* binds the bi-directional promoter between *ctcS* and the divergently encoded *ctrR*, which encodes the pathway efflux pump. Binding of CtcS to this promoter is attenuated by chlortetracycline, thereby inducing transcription of *ctrR* ^23^.

Redox-sensitive MarR-family regulators have been reported previously but these are all associated with the regulation of genes involved in oxidative stress resistance and virulence rather than antibiotic biosynthesis. In *Mycobacterium tuberculosis*, the redox-sensing HypS, regulates genes involved in antibiotic resistance in response to accumulating oxidizers like HOCl. Upon oxidation of the cysteine residues in each HypS monomer, inter-subunit disulphides form to generate a cross-linked dimer that is no longer able to bind DNA, resulting in derepression of gene expression ^17^. However, ForJ appears to behave in the opposite manner; oxidation of the cysteine residue seems to increase the strength of DNA binding, as shown by the increased non-sequence specific binding of oxidized ForJ to DNA *in vitro* and the decreased expression of *for* BGC transcripts in cultures exposed to diamide. This is more reminiscent of the MarR-family regulator BifR in *Burkholderia thailandensis*. BifR regulates genes involved in biofilm formation, particularly *escC*, which encodes a putative LasA protease. Repression of *escC* by BifR is stronger under oxidizing conditions due to the formation of disulphide-linked dimer-of-dimers upon oxidation of a cysteine residue located within the DNA-binding domain ^18^.

To our knowledge this is the first report of a redox sensitive MarR-family regulator controlling antibiotic biosynthesis in bacteria in response to oxidizing conditions. However, many *Streptomyces* species and other antibiotic-producing bacteria have lowered antibiotic production during liquid culture, where the levels of oxygen are high, and this is a significant barrier to antibiotic discovery and development, particularly because isolating clinically useful amounts of these molecules often requires large-scale liquid fermentation. Understanding how this redox regulation works is the first step towards engineering bacteria to produce such antibiotics in liquid culture. Based on the paradigm of ForJ and formicamycin biosynthesis, we propose that targeting redox-sensitive regulatory genes in other antibiotic biosynthesis pathways could provide a simple way to switch on antibiotic biosynthesis in solid and liquid cultures.

## Acknowledgments

This work was supported by the BBSRC via the Institute Strategic Programme Grant ‘Harnessing Biosynthesis for Sustainable Food and Health’ (HBio) (BB/X01097X/1), responsive mode grant BB/S00811X/2 to M.I.H and B.W (R.D) and Discovery Fellowship BB/X00967X/1 to R.D. K.N and H.P.M were supported by BBSRC PhD studentships (BBSRC Doctoral Training Program grant BB/M011216/1) and responsive mode grant BB/Y005724/1 to M.I.H and B.W (K.N). We thank Colin Grant at ReFeyn Ltd, Oxford, for the analysis of ForJ on the ReFeyn 2 machine at ReFeyn Ltd. Oxford in February 2022.

## Author contributions

Conceptualization, R.D, K.N, M.I.H, and B.W; Methodology, R.D, K.N, C.S and E.S.H; Formal analysis R.D, K.N, C.S, C.O.M, G.S, H.P.M and E.S.H; Investigation, R.D, K.N, C.S, C.O.M, G.S, H.M and E.S.H; Writing – Original Draft, R.D., K.N, M.I.H and B.W, Writing – Review & Editing, all authors; Visualisation, R.D and K.N; Supervision, M.I.H and B.W; Project administration, R.D, K.N, M.I.H and B.W; Funding acquisition, R.D, M.I.H and B.W.

## Data availability

The mass spectrometry proteomics data have been deposited to the ProteomeXchange Consortium via the PRIDE [1] partner repository with the dataset identifier PXD061171 with reviewer username and password pRb69151fPT0 ^24^.

## Methods

Unless otherwise stated chemicals and reagents used were of laboratory grade or above and purchased from Sigma Aldrich (UK), Thermo Fisher Scientific (UK) or Merck (UK), and enzymes were purchased from New England Biolabs.

## Generating *S. formicae* mutants

### PCR amplification of DNA fragments

DNA fragments of interest were amplified using Q5 DNA polymerase according to the manufacturer’s instructions. The following conditions were used in the thermocycler:

1. 95 °C, 2 min for initial denaturation
2. 30 cycles of:

a. 95 °C, 30 sec, denaturation
b. 55-72 °C, 30 sec, primer annealing
c. 72 °C, 30 sec per kb, extension
3. 72 °C, 10 min, for final extension
4. 4 °C final hold

PCR products were purified using gel electrophoresis. Gels were made with 1% agarose in TBE buffer (90 mM Tris HCl, 90 mM Boric Acid, 2 mM EDTA) containing 2 μg/mL ethidium bromide. DNA samples and loading buffer (5x) (0.25% (w/v) bromophenol blue, 0.25% (w/v) xylene-cyanol blue, 40% (w/v) sucrose in water) were run alongside a 1 kb plus DNA ladder plus loading dye for size determination. Electrophoresis occurs at 120 V (Sub-Cell GT electrophoresis system, BIOLINE) for 30-60 min depending on size (larger fragments were run longer for clearer separation). DNA was visualised by UV-light using a Molecular Imager Gel Doc System (BIO-RAD). Gel fragments containing DNA bands of interest were excised using a scalpel and extracted using a QiaQuick Gel Extraction Kit (QIAGEN), according to the manufacturer’s instructions.

### Plasmid preparation

Plasmid DNA was prepared using QIAprep Spin Miniprep kits (QIAGEN) from 3-5 mL overnight cultures as per manufacturer’s instructions.

### Restriction digest

Restriction enzymes were used to digest plasmid DNA and PCR fragments in 50 µL total volumes in accordance with manufacturer’s guidelines in CutSmart Buffer. After heat inactivation of the restriction enzymes, 2 μL shrimp alkaline phosphatase was added to dephosphorylate the digested plasmid DNA to prevent re-ligation. Digests were analysed by gel electrophoresis and the desired bands excised for downstream applications.

### Golden Gate assembly

gRNA sequences were ordered as single stranded oligos with BbsI overhangs from Integrated DNA technologies (IDT) and annealed at equal molarity in HEPES buffer by heating at 95 °C for 5 min before cooling to 4 °C (at 1 °C/s). To assemble into the pCRISPomyces-2 vector, 20 μL reactions with 100 ng purified backbone and 0.3 μL insert in the presence of 2 μL T4 ligase buffer and 1 μL T4 ligase with 1 μL BbsI and dH_2_O were set up. This was incubated in a thermocycler under the following conditions:

- 10 cycles of the following:

`o` 10 min at 37 °C
`o` 10 min at 16 °C
- 5 min at 50 °C
- 20 min at 65 °C
- 4 °C hold

To confirm insertion of the gRNA, the Golden Gate reaction was transformed into *E. coli* and plated on LB agar containing 100 mM IPTG and 50 μg/ml X-Gal for blue-white screening. Plasmids were subsequently purified and sequenced by Eurofins.

### Gibson assembly

Multiple DNA fragments were assembled into digested vector backbones using designed overlaps of between 18 and 24 nucleotides. Gel extracted DNA fragments were incubated in a ratio of 1:3 of plasmid to insert (1:5 for inserts smaller than 300 nucleotides) in the presence of Gibson Assembly master mix at 50 °C for 1 h.

### Transformation of competent *E. coli*

For transformation, approximately 2 μg of DNA was added to 50 μl of cells. For electrocompetent cells, the mixture was transferred to an ice-cold electroporation cuvette and electroporation was carried out using the BioRad Electroporator set to 200 Ω, 25 μF and 2.5 kV. For chemically competent cells, the mixture was incubated on ice for around 30 min, before being heat-shocked for 30 s at 42 °C then immediately cooled on ice for 2 min. The transformed cells were diluted in LB and recovered at 37 °C for 1 h with 220 rpm shaking before plating on selective media for overnight incubation and colony selection. After overnight incubation, single colonies were picked and tested by colony PCR using BIOTAQ Red DNA Polymerase (Bioline) according to the manufacturer’s instructions.

### Conjugation into *S. formicae*

Single colonies of non-methylating *E. coli* ET12567/pUZ8002 containing the required plasmid were selected from plates and grown in 10 mL LB broth plus antibiotics at 37 °C overnight at 220 rpm. Subcultures of OD_600_ between 0.4 and 0.6 were washed twice in LB to remove antibiotics. Either 50 μL or 200 μLof a *S. formicae* spore suspension was heat shocked at 50 °C for 10 min in 500 μL 2YT to encourage germination and added to the washed *E. coli* cells (more spores used for CRISPR deletions, less for integrative vectors). The cell mixture was pelleted by centrifugation at 15,871 xg for 1 min, the supernatant was removed, and cells were resuspended in the residual liquid. This was plated onto soya flour mannitol (SFM) agar containing 10 mM MgCl_2_ and incubated at 30 °C for 16-20 h. For selection of desired ex-conjugants, 0.5 mg Nalidixic acid and an appropriate concentration of the selection antibiotic (to give final selective concentration in the plate) was added in 1 mL dH_2_O to each plate and cultures returned to the 30°C incubator for 5 days or until colonies appeared. Following purification by re-streaking on antibiotic selective media, *Streptomyces* ex-conjugants were confirmed by colony PCR. Single colonies were picked and soaked in 100 μL 50% DMSO at 50 °C for 1 h. This was then used as template for PCR using PCRBIO Taq Mix Red (PCR Biosystems) at 10% of the final volume of the reaction (usually 2.5 μL in 25 μL). For pCRISPomyes-2 edited strains, colonies that were confirmed to contain the desired mutation were then re-streaked for multiple generations on non-selective media to encourage the loss of the plasmid.

## Titre analysis of *S. formicae* mutants

*S. formicae* WT and mutant strains were grown as confluent lawns on SFM agar at 30 °C for 10 days. Equal size agar plugs (1 cm^3^) were taken in triplicate from each plate and each one shaken with ethyl acetate (1 mL) for 1 h before being centrifuged at 3000 xg for 5 min. An aliquot of the ethyl acetate extract (300 μL) was transferred to a clean tube and the solvent was removed under reduced pressure. The resulting extract was dissolved in methanol (200 μL) before being analysed by HPLC (Agilent 1290 UHPLC). For liquid cultures, single colonies were used to inoculate 10 mL seed cultures in TSB media and grown at 30 °C, 200 rpm for 2 days. Seed cultures were diluted 1:100 into 10 mL liquid SFM and grown at 30 °C, 200 rpm for 10 days. A 1 mL sample was taken and shaken with an equal volume of ethyl acetate for 1 h at room temperature. An aliquot of the resulting ethyl acetate (300 μL) was transferred to a clean tube, the solvent removed under reduced pressure, and the resulting extract resuspended in methanol (200 μL) for analysis. Formicamycin and fasamycin production titres were quantified by HPLC on an Agilent 1290 system fitted with a Phenomenex Kinetex 5 μm XB-C_18_ 100 Å column (100 mm × 4.1 mm) using the following gradient: solvent A: water with 0.1 % v/v formic acid, solvent B: acetonitrile with 0.1 % v/v formic acid; flow rate: 1 mL/min; injection volume: 10 μL; *T* = 0 min, 50% B; *T* = 0.5 min, 50% B; *T* = 12 min, 98% B; *T* = 13 min, 98% B; *T* = 13.5 min, 50% B; *T* = 15 min, 50% B; UV monitoring at 285 nm (formicamycins) and 418 nm (fasamycins). Peak area integration was performed using OpenlabCDS ChemStation software and titres determined by comparison with calibration curves. Calibration curves were produced using the same HPLC method and solutions of formicamycin J (10, 20, 40, 100, 200, 400, 800 μM) and fasamycin L (1, 2.5, 5, 10, 20, 50 μM) (**Fig. S5** and **S6**).

## Protein production and purification

Protein production plasmids were transformed into fresh *E. coli* and grown at 37 °C for 3 h and induced overnight with 0.1 mM IPTG at 18 °C, 200 rpm shaking. The following day, cells were harvested by centrifugation at 20,600 xg for 20 min. The pellets were flash frozen in liquid nitrogen and either used immediately or stored at -80 °C for purification at a later date. Pellets were thawed on ice and resuspended in lysis buffer (20 mM Tris, pH 7.5, 75 mM NaCl, 1 mM TCEP, 0.1% Triton-X, 1 mg/mL DNase, 1 mg/mL Lysozyme, cOmplete^TM^ EDTA-free protease inhibitor cocktail), sonicated until and centrifuged at 45,000 xg for 30 min. A His-trap HP (5mL) column was loaded onto an AKTA FPLC (AKTA pure) and washed with 20% ethanol before equilibrating with wash buffer (80 mM Tris, pH 7.5, 200 mM NaCl, 1 mM TCEP, 20 nM Imidazole). Lysate was applied to the column through the sample line at 5 mL/min and the column washed with 20 volumes of wash buffer. Protein was eluted in elution buffer (80 mM Tris, pH 7.5, 200 mM NaCl, 1 mM TCEP, 500 nM Imidazole) in 1 mL fractions and the presence of protein was monitored by UV absorption at 280 nm. For gel filtration, the column was equilibrated overnight into gel filtration buffer (50 mM Tris, 100 mM NaCl, 1mM TCEP, pH 7.5) and the sample was applied from the sample line (<5 mL volume). Eluted fractions were analysed by SDS-Page gel electrophoresis by mixing with bromophenol blye dye (+ β-mercaptoethanol), boiling (100 °C, 10 min) and loading into Invitrogen Novex^TM^ Tris-Glycine Gels (Thermo Fisher) gels with Novex^TM^ Tris-Glycin SDS Running Buffer. Samples were run at 150V for 60 min alongside Colour Protein Standard (NEB). Gels were stained with InstantBlue Protein Stain (Expedeon) with agitation at room temperature for 1 h or overnight. Gels were de-stained in dH_2_O for at least 1 h and imaged using white light on a Molecular Imager Gel Doc System (BIORAD). Protein samples were concentrated in Amicon centrifuge spin filter tubes at 4 °C, 3130 xg and concentrations were assessed using Nanodrop and Qubit protein broad range assay according to the manufacturer’s instructions.

## ReDCaT SPR

ReDCaT SPR was conducted as previously reported (Stevenson et al., 2013). Briefly, single stranded biotinylated DNA-linker sequence (gcaggaggacgtagggtagg) was applied to a SA Chip (Cytiva) and immobilised using the Biocore 8K+ SPR system. In general, SPR experiments were run in HBS-EP+ buffer and water. A solution of 1M NaCl/50 mM NaOH was used to regenerate the chip after each cycle. Double stranded oligos for promoters of interest were prepared by annealing single stranded oligos at equal molarity by heating at 90 °C for 2 min and ramping to 4°C at 0.1 °C/sec in a thermocycler. The annealed oligos were diluted to 1000 nM in HBS-EP + buffer and stored for use. Protein binding was tested at three concentrations, usually 1000, 100 and 10 nM. Each cycle consisted of a 60 s capture stage at 10 μL/min where DNA is captured to the chip, a 60 s analyte binding stage with 60 secs dissociation at 50 μL/min for protein to bind and stabilise, and a 60 s regeneration step at 10 μL/min to remove protein and DNA probes for the following cycle. Binding responses were recorded at both early and late time-points within the analyte binding stage, but late binding was used to calculate %Rmax as a representation of stable binding.

## Western blot analysis of 6xHis tagged proteins in *S. formicae*

Tagged proteins were expressed *in trans* on integrative vectors. Strains were grown on SFM agar for 2-3 days, before mycelium was harvested with a sterile spatula. Samples were subjected to SDS-PAGE as described in protein production and purification, before being transferred to a nitrocellulose Biodyne A membrane (Pall Corporation) in a Trans-Blot SD Semi-Dry Transfer Cell (BIORAD). Three layers of blotting paper, equal size to the gel, were soaked in 1 x transfer buffer (25 mM Tris, 192 mM Glycine, 0.1 % SDS + 20 % methanol) and placed on the transfer cell anode plate. Nitrocellulose membrane was soaked in methanol (1 min), followed by washing in transfer buffer (5 min) and placed on top of the blotting paper. The SDS polyacrylamide gel of proteins to be transferred was placed on top of the membrane, followed by three more layers of soaked blotting paper before transfer took place (10 V, 1 h). A blocking solution of 5% (w/v) fat-free skimmed milk powder in 1 x TBST (50 mM Tris Cl pH 7.5, 150 mM NaCl, 1% Tween) was poured over the membrane and incubated at room temperature overnight with gentle agitation. Anti-Flag antibody conjugated to horseradish peroxidase (HRP) was diluted 1:20,000 in 1 x TBST and used to check strains containing 3xFlag tagged proteins. The membrane was incubated in 20 mL of the respective antibody suspension at room temperature for 1 hr and washed 3 times for 10 min in TBST. Membranes were developed for 1 min in a 50:50 mix of solutions A and B and fluorescence was detected using the ECL setting for imaging using a SYNGENE G:Box.

| **Developing Solution A** | **Developing Solution B** |
| --- | --- |
| 10 mL 100 mM Tris pH 8.5 | 10 mL 100 mM Tris pH 8.5 |
| 0.01% luminol | 0.00018 % hydrogen peroxide |
| 0.0045 % coumaric acid |  |

## Gene expression analysis of *S. formicae* mutants

For all RNA work, RNase-free water was prepared by DEPC treating (0.1% v/v) at 37 °C O/N and autoclaving twice before use. All other equipment required for the processing of RNA was double autoclaved before use. Samples from liquid culture were harvested by centrifugation, and solid cultures were harvested using a sterile spatula from plates with cellophanes. Samples were flash frozen in liquid nitrogen for storage at -80 °C before extraction. Pellets were thawed and crushed in liquid nitrogen using a sterile pestle and mortar on dry ice. Samples were resuspended in 1 mL RLT Buffer (Qiagen) supplemented with β-mercaptoethanol (10 mL in every 1 mL buffer) and vortexed for 1 min. Samples were then applied to a QIA-shredder column (Qiagen) and centrifuged at 15,871 xg for 2 min. Flow through was collected (leaving the pellet behind) and mixed with 700 mL acid-phenol:chloroform, pH 4.5 (with IAA, 125:24:1) for 1 min. Samples were then incubated at room temperature for 3 min before centrifuging at 15,871 xg for 20 min. The upper phase was collected and mixed with 0.5 volumes of 95% ethanol. This was then applied to a RNeasy Mini spin column (Qiagen) and purified following the manufacturers protocol including on column DNase treatment. Following elution, the Turbo-DNase kit was then used according to the manufacturers protocol and a further Qiagen RNeasey mini clean-up was conducted. Samples were then aliquoted for quantification or other downstream processes and flash frozen in liquid nitrogen for storage at –80 °C. For quantification, RNA was analysed by Nanodrop.

RNA samples were confirmed to be DNA-free by conducting test PCRs on the 16S rRNA gene and converted to cDNA using the LunaScript RT SuperMix Kit (NEB) according to the manufacturer’s instructions. Primers for each transcript were optimised using serial dilutions of ePAC template DNA and checked for specificity by melt-curve analysis and gel electrophoresis. Reactions were run in biological triplicate and technical duplicate using the Luna Universal qPCR Master Mix according to the manufacturer’s guidelines in a total volume of 20 μL with approximately 100 ng template cDNA and a final concentration of 0.25 μM of each primer. ΔCT values were normalised to the 16S rRNA gene.

## Quantitative proteomics using isobaric Tandem Mass Tag (TMT) labelling

For proteomics, strains were grown on SFM agar for 5 days (on cellophanes) and the biomass scraped into falcon tubes using a sterile spatula. Pellets were resuspended in lysis buffer (2% SDS, 50 mM TEAB Buffer pH 8.0, 150 mM NaCl, cOmplete^TM^ EDTA-free protease inhibitor cocktail). Cells were homogenised 3 times and heated for 10 min in a boiling water bath. Samples were sonicated 4 times for 20 s each cycle, with 1 min between each cycle and centrifuged at 3130 xg for 30 min at room temperature to eliminate cell debris. Protein concentration was assayed using Qubit according to the manufacturer’s instructions and 1 mg was removed into a fresh 15 mL falcon. Proteins were precipitated with chloroform/methanol ^26^.

The pellets were washed with acetone and dissolved in 100 µL of 0.2M EPPS buffer pH 8 with 2.5% sodium deoxycholate (SDC) and vortexed under heating in a boiling water bath. Protein concentration was determined using a BCA assay and 100 µg protein from each sample was used. Cysteine residues were reduced with dithiothreitol, alkylated with iodoacetamide, and the proteins digested with Sequencing Grade Modified Trypsin (Promega) in the SDC buffer according to the manufacturer. After digestion, the SDC was precipitated by adjusting to 0.2% TFA, and the clarified supernatant subjected to C_18_ solid phase extraction (SPE; OMIX tips; Agilent). TMT labelling was performed using a Thermo TMT16plex kit (lot number: VK309613) according to manufacturer’s instructions with slight modifications, as follows; the dried peptides were dissolved in 90 µL of 0.2 M EPPS buffer pH 8, 10 % acetonitrile, and 250 µg TMT16plex reagent dissolved in 22 µL acetonitrile was added. After the labelling, aliquots of 1.7 µL from each sample were combined and an aliquot of the mix was analysed by LCMS (methods see below) to check labelling efficiency and estimate total sample abundances. The main sample aliquots were then combined correspondingly to achieve equal level of peptides and desalted using a 50 mg C_18_ Sep-Pak cartridge (Waters). The eluted peptides were dissolved in 500 µL of 25 mM NH_4_HCO_3_ and fractionated by high pH reversed phase HPLC. Using an ACQUITY Arc Bio System (Waters), the samples were loaded to a Kinetex 5 µM EVO C_18_ 100 Å LC Column 250×4.6 mm (Phenomenex). Fractionation was performed with the following linear (unless stated otherwise) steps of a gradient of solvents A (water), B (acetonitrile), and C (25 mM NH_4_HCO_3_) at a flow rate of 1 mL/min: solvent C was kept at 10% throughout the gradient; *T* = 0 min, 5% B; *T* = 5 min, 5% B; *T* = 10 min, 10% B (curve 4); *T* = 60 min, 40% B; *T* = 75 min, 80% B; *T* = 78 min, 80% B; *T* = 79 min, 5% B; *T* = 105 min, 5% B. Fractions of 1 mL were collected, dried down, and concatenated to produce 21 final fractions for LCMS analysis.

Aliquots of all concatenated fractions were analysed by nanoLC-MS/MS on an Orbitrap Eclipse Tribrid mass spectrometer coupled to an UltiMate 3000 RSLCnano LC system (Thermo Fisher Scientific). The samples were loaded onto a trap column (nanoEase M/Z Symmetry C_18_ Trap Column, Waters) with 0.1% v/v TFA at 15 µL/min for 3 min. The trap column was then switched in-line with the analytical column (nanoEase M/Z column, HSS C_18_ T3, 1.8 µM, 100 Å, 250 mm × 0.75 µm, Waters) for separation using the following gradient of solvents A (water + 0.1 % v/v formic acid) and B (80:20 acetonitrile/water + 0.1% v/v formic acid) at a flow rate of 0.2 µL/min: *T* = 0 min, 3% B; *T* = 3 min, 3% B; *T* = 10 min, 8% B (curve 4); *T* = 80 min, 35% B; *T* = 105 min, 53 % B; *T* = 113 min, 99% B; *T* = 116 min, 99% B; *T* = 117 min, 3% B; *T* = 140 min, 3% B.

Mass spectrometry data were acquired with the following parameters in positive ion mode: MS1/OT: resolution 120K, profile mode, mass range m/z 400-1800, AGC target 4e5, max. inject time 50 ms; MS2/IT: data-dependent analysis with the following parameters: IT rapid mode, centroid mode, quadrupole isolation window 0.7 Da, charge states 2-5, threshold 1.9e4, CID fragmentation with CE=30, AGC 1e4, max. inject time 50 ms, dynamic exclusion one count/15 s/±7 ppm; MS3 synchronous precursor selection (SPS): 10 SPS precursors, isolation window 0.7 Da, HCD fragmentation with CE=50, Orbitrap Turbo TMT and TMTpro resolution 30 k, AGC target 200 %, max inject time 105 ms, Real Time Search: protein database *S. formicae* (uniprot.org, 8132 entries), enzyme trypsin, 1 missed cleavage, oxidation (M) as variable, carbamidomethyl (C) and TMTpro as fixed modifications, Xcorr=1, dCn=0.05.

The acquired raw data were processed and quantified in Proteome Discoverer 3.2 (Thermo Fisher Scientific); all mentioned tools of the following workflow are nodes of the proprietary Proteome Discoverer (PD) software.

The *S. formicae* fasta database (uniprot.org, 8132 entries) was imported into PD adding a reversed sequence database for decoy searches; a database for common contaminants (maxquant.org, 245 entries) was also included. The database search was performed using the incorporated search engines CHIMERYS (MSAID, Munich, Germany) and Comet ^27^.

The processing workflow included recalibration (RC) of MS1 spectra, reporter ion quantification by most confident centroid (20 ppm) and a search on the imported *S. formicae* database. The Top N Peak Filter was applied with 20 peaks per 100 Da. For CHIMERYS, the inferys_4.7.0_fragmentation prediction model was used with fragment tolerance of 0.3 Da, enzyme trypsin with 1 missed cleavage, variable modification oxidation (M), fixed modifications carbamidomethyl (C) and TMTpro on N-terminus and K. For Comet the version 2019.01 rev.0 parameter file was used with default settings except precursor tolerance set to 5 ppm and trypsin missed cleavages set to 1. Modifications were the same as for CHIMERYS.

The consensus workflow included the following parameters: intensity-based abundance, normalisation on total peptide abundances, protein abundance-based ratio calculation, only unique peptides (protein groups) for quantification, TMT channel correction values applied (Lot VK309613), co-isolation/SPS matches/CHIMERYS Coefficient thresholds 50%/65%, 0.8, missing values imputation by low abundance resampling, hypothesis testing by t-test (background based), adjusted p-value calculation by BH-method. The results were exported into a Microsoft Excel table including data for normalised and un-normalised abundances, ratios for the specified conditions, the corresponding p-values and adjusted p-values, number of unique peptides, q-values, PEP-values, identification scores from both search engines; FDR confidence filtered for high confidence (strict FDR 0.01) only.

## Production and purification of Formicamycin I and Fasamycin E

Wildtype or Δ*forX S. formicae* were used for the production of formicamycin I and fasamycin E, respectively as previously reported ^9,20^.

