## Supplementary Information for "Redox control of antibiotic biosynthesis"

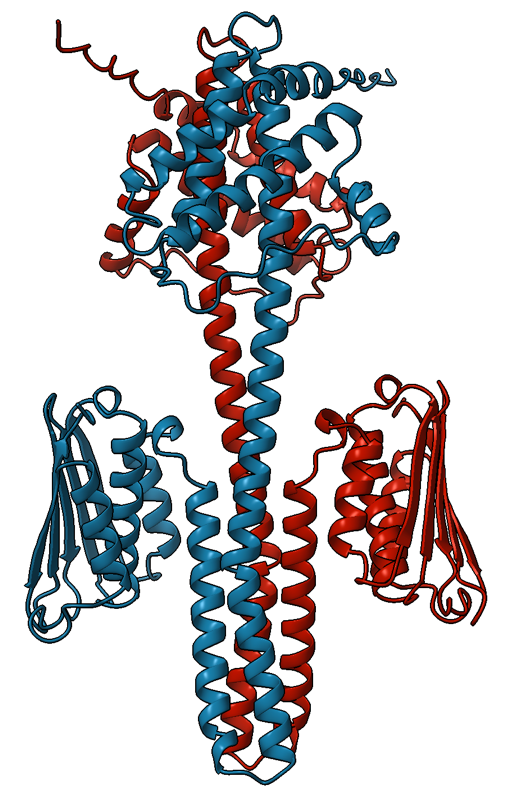

**Figure S1.** AlphaFold3 predicted structure of ForG dimer

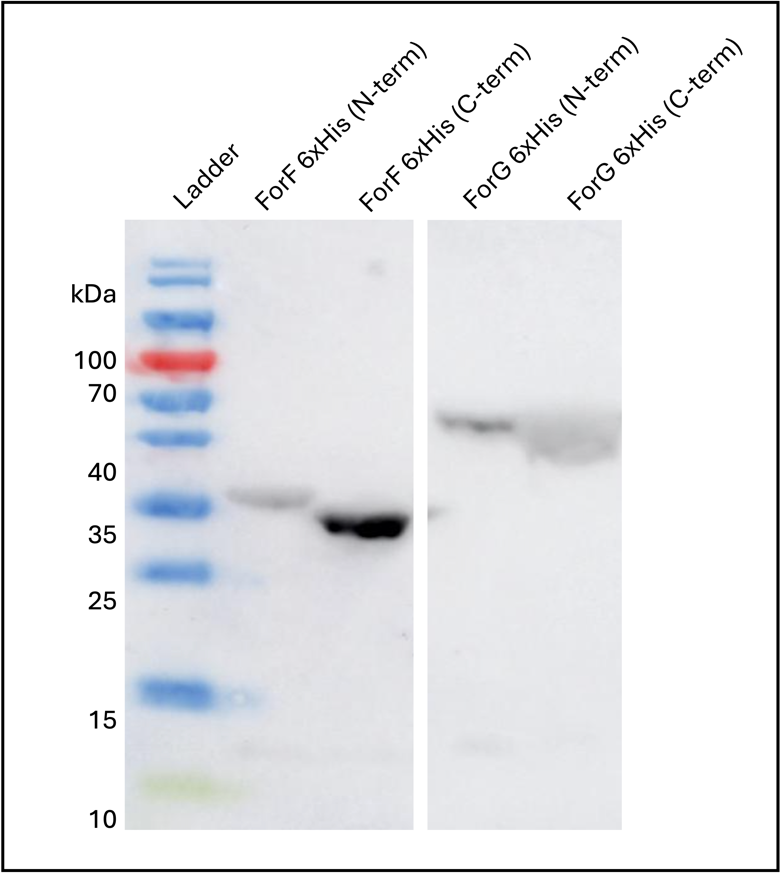

**Figure S2.** Protein purification prep of ForG from *E. coli* Nico21 using the pET28a(+) and pET29a(+) vectors

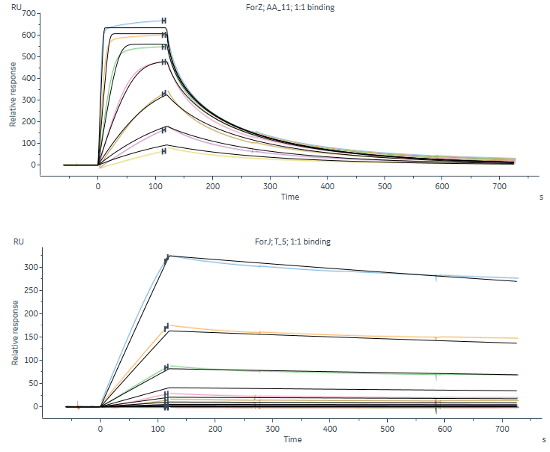

**Figure S3.** Multi-cycle affinity and kinetics for ForZ against DNA probe AA_11 and ForJ against DNA probe T_5. Experiments were run in duplicate, using standard SPR buffer (see methods section ‘ReDCaT SPR’) with the addition of 1mM DTT to ForJ experiments. ForZ binds DNA with a K_D_ of 46 nM whereas ForJ binds more weakly, with a K_D_ of 288 nM.

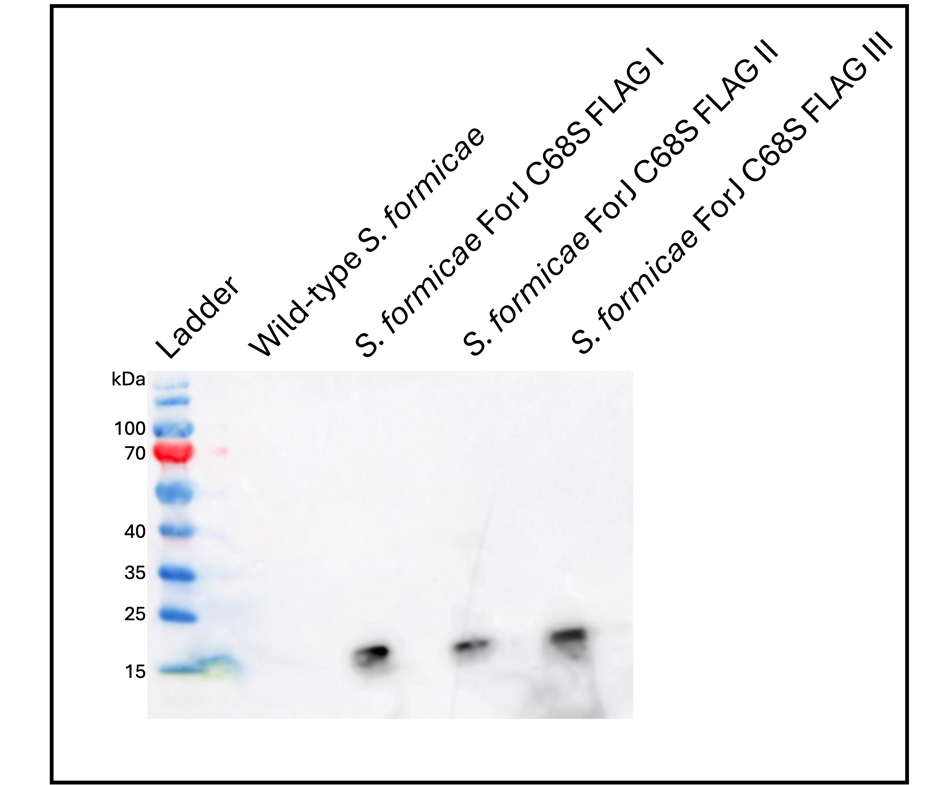

**Figure S4.** Western blot analysis of S. formicae producing ForJ C68S 3x-Flag (n=3) suggests C68 is not essential for protein folding and stability *in vivo*

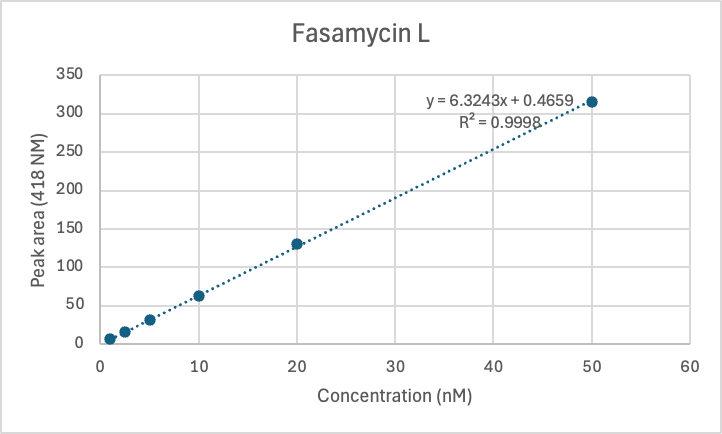

**Figure S5.** Calibration curve for Fasamycin L (average of 3 replicates)

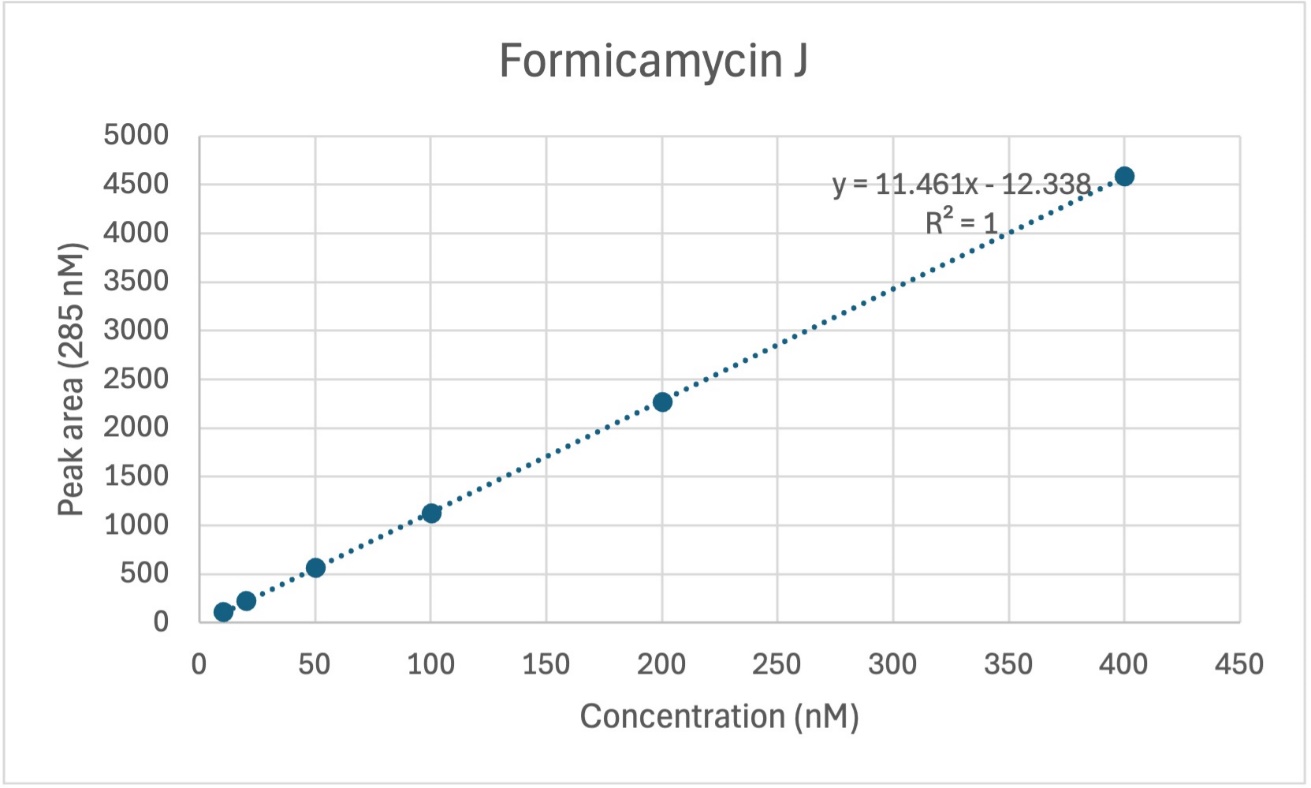

**Figure S6.** Calibration curve for Formicamycin J (average of 3 replicates)

**Strains used and generated in this study:**

| **Strain** | **Description/Genotype** | **Plasmid/Resistance** | **Reference/Source** |
| --- | --- | --- | --- |
| E. coli ET12567 | dam- dcm- hsdS- | pUZ8002, Cml^R^/Tet^R^ | (MacNeil et al., 1992) |
| E. coli NEB5α | fhuA2 Δ(argF-lacZ)U169 phoA glnV44 Φ80 Δ(lacZ) M15 gyrA96 recA1 relA1 endA1 thi-1 hsdR17 |  | New England Biosciences |
| E. coli BL21 | fhuA2 [lon] ompT gal (λ DE3) [dcm] ΔhsdSλ DE3 = λsBamHIo ΔEcoRI-B int::(lacI::PlacUV5::T7 gene1) i21 Δnin5 |  | Invitrogen |
| E. coli Nico21 | BL21 derivative, can:: CBD fhuA2 [lon] ompT gal (λ DE3) [dcm] arnA::DB sly::CBD glmS6Ala Δ ΔhsdSλ DE3 = λsBamHIo ΔEcoRI-B int::(lacI::PlacUV5::T7 gene1) i21 Δnin5 |  | Invitrogen |
| E. coli C43 | BL21 derivative, F – ompT hsdSB (rB- mB-) gal dcm (DE3) |  | Sigma |
| E. coli Nico21 pET29a(+) ForF |  | pET29a(+) ForF/ Kan^R^ | This work |
| E. coli BL21 pET28a(+) ForZ |  | pET28a(+) ForZ/ Kan^R^ | This work |
| E. coli C43 pET28a(+) ForJ |  | pET28a(+) ForJ/ Kan^R^ | This work |
| S. formicae wild- type |  |  | Lab stock |
| S. formicae ΔforGF |  |  | Devine et al., 2021 |
| S. formicae ΔforGF::forGF |  |  | Devine et al., 2021 |
| S. formicae ΔforG |  |  | This work |
| S. formicae ΔforF |  |  | This work |
| S. formicae ForF D53E |  |  | This work |
| S. formicae ForF D53A |  |  | This work |
| S. formicae ΔforF::forF D53E |  | pMS82 ForF D53E /Hyg^R^ | This work |
| S. formicae ΔforF::forF D53A |  | pMS82 ForF D53E /Hyg^R^ | This work |
| S. formicae ΔforJ |  |  | Devine et al., 2021 |
| S. formicae ForJ-3xFlag |  | pMS82 ForJ-3xFlag/Hyg^R^ | Devine et al., 2021 |
| S. formicae ForJ C68S |  |  | This work |
| S. formicae ForJ C68S 3xFlag |  | pMS82 ForJ C68S-3xFlag/Hyg^R^ | This work |

**Plasmids used and generated in this study:**

| **Plasmid** | **Description** | **Resistance** | **Reference/Source** |
| --- | --- | --- | --- |
| pUZ8002 | RK2 derivative with a mutation in oriT | Kan^R^ | (Keiser et al., 2000) |
| pCRISPomyces-2 | oriT, _rep_pSG5(ts), ori^ColE1^, sSpcas9, synthetic guide RNA cassette | Apr^R^ | (Cobb, Wang and Zhao, 2015) |
| pET28a(+) | pBR322 origin and fI origin, expression vector | Kan^R^ | Invitrogen |
| pET29a (+) | pBR322 origin and fI origin, expression vector | Kan^R^ | Invitrogen |
| pMS82 | ori, pUC18, hyg, oriT, RK2, int ΦBT1 | Hyg^R^ | (Gregory, Till and Smith, 2003) |
| pET28a(+) ForG | Codon optimised ForG (GenScript) cloned between NdeI/EcoRI | Kan^R^ | This work |
| pET29a(+) ForF | Codon optimised ForF (GenScript) cloned between NdeI/EcoRI | Kan^R^ | This work |
| pET28a(+) ForZ | Native ForZ cloned between NdeI/EcoRI | Kan^R^ | This work |
| pET28a(+) ForJ | Codon optimised ForJ (GenScript) cloned between NdeI/EcoRI | Kan^R^ | This work |
| pCRISPomyces-2 ΔforG | For the deletion of forG | Apr^R^ | This work |
| pCRISPomyces-2 ΔforF | For the deletion of forF | Apr^R^ | This work |
| pCRISPomyces-2 ForF D53E | For the substitution of ForF D53 to E53 | Apr^R^ | This work |
| pCRISPomyces-2 ForF D53A | For the substitution of ForF D53 to A53 | Apr^R^ | This work |
| pCRISPomyces-2 ForJ C68S | For the substitution of ForJ C68 to S68 | AprR | This work |
| pMS82 ForJ C68S 3x Flag | For in trans expression of ForJ C68S 3x-Flag (synthesised by GenScript) in a forJ deletion background | HygR | This work |

**Primers used and generated in this study:**

| **Primer** | **Description** | **Sequence (5’-3’)** |
| --- | --- | --- |
| pET28/29a Test F | To test inserts into the NdeI/EcoRI site of pET28a | TAATACGACTCACTATAGGG |
| pET28/29a Test R | To test inserts into the NdeI/EcoRI site of pET28a | TCGCCACCTCTGACTTGAGCGTCGA |
| pCRISPomyces-2 Test F | To test inserts into the XbaI site of pCRISPomyes-2 | AGGCTAGTCCGTTATCAACTTGAAA |
| pCRISPomyces-2 Test R | To test inserts into the XbaI site of pCRISPomyes-2 | TCGCCACCTCTGACTTGAGCGTCGA |
| gRNA test | To sequence across the BbsI site of pCRISPomyces-2 | ATACGGCTGCCAGATAAGGC |
| ForZ pET28a F | To amplify ForZ for assembly into the NdeI/EcoRI site of pET28a by Gibson assembly | CCGCGCGGCAGCCATATGGCTAGCggacttcaagccatggcctcc |
| ForZ pET28a R | To amplify ForZ for assembly into the NdeI/EcoRI site of pET28a by Gibson assembly | AGCTTGTCGACGGAGCTCGAATTCtcaccgctcgcacgccgc |
| ForF pET29a F | To amplify ForF for assembly into the NdeI/EcoRI site of pET29a by Gibson assembly | CTTTAAGAAGGAGATATACATATGATGCAGACCGTGGTTACCCG |
| ForF pET29a R | To amplify ForF for assembly into the NdeI/EcoRI site of pET28a by Gibson assembly | GTGCTCGAGTGCGGCCGCAAGCTTACCACGATCACCTTGCGGC |
| ForG pET29a F | To amplify ForG for assembly into the NdeI/EcoRI site of pET28a by Gibson assembly | CTTTAAGAAGGAGATATACATATGATGAATGAGGCGGCGACCG |
| ForG pET29a R | To amplify ForG for assembly into the NdeI/EcoRI site of pET28a by Gibson assembly | GTGCTCGAGTGCGGCCGCAAGCTTACCGCTTTTGGTCGGCAGC |
| ForF gRNA F | gRNA Forward with BbsI overhangs | ACGCtggcgaagatgttgcgcaga |
| ForF gRNA R | gRNA Reverse with BbsI overhangs | AAACtctgcgcaacatcttcgcca |
| ForF KO Repair F1 | Amplify ForF flank for Gibson assembly into XbaI site of pCRISPomyces-2 | gctcggttgccgccgggcgttttttaTCTAGAaccactacgaacagcgcctg |
| ForF KO Repair R1 | Amplify ForF flank for Gibson assembly into XbaI site of pCRISPomyces-2 omitting ForF coding region | GCTGCTGCGACCAGGCGAGCTCGCcatccgcgcctcatccgctc |
| ForF KO Repair F2 | Amplify ForF flank for Gibson assembly into XbaI site of pCRISPomyces-2 omitting ForF coding region | GCGAGCTCGCCTGGTCGCAGCAGCtgaggctcaggcgggttcga |
| ForF KO Repair R2 | Amplify ForF flank for Gibson assembly into XbaI site of pCRISPomyces-2 | gcaacgcggcctttttacggttcctggccTCTAGAatcgtcgccgagctggagaa |
| ForF D53E F1 | Amplify ForF flank for Gibson assembly into XbaI site of pCRISPomyces-2 | gctcggttgccgccgggcgttttttaTCTAGAagcatcatgtggcggttcca |
| ForF D53E R1 | Amplify ForF with silent mutation in PAM | CCGACGTCGAGCAGCACCACTTCGGGCGACTTGGCCGCGATCA |
| ForF D53E F2 | Amplify ForF with silent mutation in PAM | TGATCGCGGCCAAGTCGCCCGAAGTGGTGCTGCTCGACGTCGG |
| ForF D53E R2 | Amplify ForF and change D53 to E53 | AACTtggcgaagatgttgcgcagaTGATGCTTCACGGTGCCCT |
| ForF D53E F3 | Amplify ForF and change D53 to E53 | AGGGCACCGTGAAGCATCAtctgcgcaacatcttcgccaAGTT |
| ForF D53E R3 | Amplify ForF flank for Gibson assembly into XbaI site of pCRISPomyces-2 | gcaacgcggcctttttacggttcctggccTCTAGAactggacgtacgaggaggtg |
| ForF D53A F1 | Amplify ForF flank for Gibson assembly into XbaI site of pCRISPomyces-2 | gctcggttgccgccgggcgttttttaTCTAGAagcatcatgtggcggttcca |
| ForF D53A R1 | Amplify ForF with silent mutation in PAM | CCGACGTCGAGCAGCACCACACGGGGCGACTTGGCCGCGATCA |
| ForF D53A F2 | Amplify ForF with silent mutation in PAM | TGATCGCGGCCAAGTCGCCCCGTGTGGTGCTGCTCGACGTCGG |
| ForF D53A R2 | Amplify ForF and change D53 to A53 | AACTtggcgaagatgttgcgcagaTGATGCTTCACGGTGCCCT |
| ForF D53A F3 | Amplify ForF and change D53 to A53 | AGGGCACCGTGAAGCATCAtctgcgcaacatcttcgccaAGTT |
| ForF D53A R3 | Amplify ForF flank for Gibson assembly into XbaI site of pCRISPomyces-2 | gcaacgcggcctttttacggttcctggccTCTAGAactggacgtacgaggaggtg |
| ForF Seq Test F1 | Test ForF deletion and point mutations | ggccggcgggagtccgcggc |
| ForF Seq Test R1 | Test ForF deletion and point mutations | gccatcgaacccgcctgagc |
| ForF Seq Test F2 | Test ForF deletion and point mutations | gtcgagttcctgctgcccacg |
| ForF Seq Test R2 | Test ForF deletion and point mutations | cctcggcatcgtcgccgagc |
| ForG gRNA F | gRNA Forward with BbsI overhangs | ACGCaggcgccagtcggcctggta |
| ForG gRNA R | gRNA Reverse with BbsI overhangs | AAACtaccaggccgactggcgcct |
| ForG KO Repair F1 | Amplify ForG flank for Gibson assembly into XbaI site of pCRISPomyces-2 | gctcggttgccgccgggcgttttttaTCTAGAgacgcggtgacgacgaggaa |
| ForG KO Repair R1 | Amplify ForG flank for Gibson assembly into XbaI site of pCRISPomyces-2 omitting ForG coding region | GCTGCTGCGACCAGGCGAGCTCGCggcagcctcgttcacagcag |
| ForG KO Repair F2 | Amplify ForG flank for Gibson assembly into XbaI site of pCRISPomyces-2 omitting ForG coding region | GCGAGCTCGCCTGGTCGCAGCAGCtgaggcgcggatgcagaccg |
| ForG KO Repair R2 | Amplify ForG flank for Gibson assembly into XbaI site of pCRISPomyces-2 | gcaacgcggcctttttacggttcctggccTCTAGAcgagcgcctggtcagactcg |
| ForG Seq Test F1 | Test ForG deletion and point mutations | cttggccatctgcatgagcg |
| ForG Seq Test R1 | Test ForG deletion and point mutations | gtcgtcgacgatcacgatgc |
| ForG Seq Test F2 | Test ForG deletion and point mutations | gagaccgcgcgaccgaaccg |
| ForG Seq Test R2 | Test ForG deletion and point mutations | tggacgagctgtgccgcagc |
| ForJ C68S gRNA F | gRNA Forward with BbsI overhangs | ACGCgggcacggtgatgcccaagt |
| ForJ C68S gRNA R | gRNA Reverse with BbsI overhangs | AAACacttgggcatcaccgtgccc |
| ForJ C68S F1 | Amplify ForJ flank for Gibson assembly into XbaI site of pCRISPomyces-2 | gctcggttgccgccgggcgttttttaTCTAGAcgtcgccatgagcgtgaccg |
| ForJ C68S R1 | Amplify ForJ with silent mutation in PAM | cggtgatgcccaagtcTgcg |
| ForJ C68S F2 | Amplify ForJ with silent mutation in PAM | cgcAgacttgggcatcaccg |
| ForJ C68S R2 | Amplify ForJ and change C68 to S68 | ggaggcatcgcTacgcagcttc |
| ForJ C68S F3 | Amplify ForJ and change C68 to S68 | gaagctgcgtAgcgatgcctcc |
| ForJ C68S R3 | Amplify ForJ flank for Gibson assembly into XbaI site of pCRISPomyces-2 | gcaacgcggcctttttacggttcctggccTCTAGAccacgtcgaggtgcaggtgc |
| ForJ C68S Seq Test F | Test ForJ C68S | gtctcgaagcacgtcacagc |
| ForJ C68S Seq Test R | Test ForJ C68S | cgtacgcggacgaactcctg |
| pforHI/GF F1 | SPR oligo for pforHI/GF promoter | GAGTCGGTTCCGGTGTTCTGCGCCGGGGACCCGCGCCCCG |
| pforHI/GF R1 | SPR oligo for pforHI/GF promoter with linker | CGGGGCGCGGGTCCCCGGCGCAGAACACCGGAACCGACTCcctaccctacgtcctcctgc |
| pforHI/GF F2 | SPR oligo for pforHI/GF promoter | GGGACCCGCGCCCCGCCCCGGCCCGGTCGGCCGGGTCGGC |
| pforHI/GF F2 | SPR oligo for pforHI/GF promoter with linker | GCCGACCCGGCCGACCGGGCCGGGGCGGGGCGCGGGTCCCcctaccctacgtcctcctgc |
| pforHI/GF F3 | SPR oligo for pforHI/GF promoter | GTCGGCCGGGTCGGCATGGTGTGGTCGGGCGCTGCTCACG |
| pforHI/GF F3 | SPR oligo for pforHI/GF promoter with linker | CGTGAGCAGCGCCCGACCACACCATGCCGACCCGGCCGACcctaccctacgtcctcctgc |
| pforHI/GF F4 | SPR oligo for pforHI/GF promoter | CGGGCGCTGCTCACGGTCATCGTGTACCCCCTGTGCACGA |
| pforHI/GF F4 | SPR oligo for pforHI/GF promoter with linker | TCGTGCACAGGGGGTACACGATGACCGTGAGCAGCGCCCGcctaccctacgtcctcctgc |
| pforHI/GF F5 | SPR oligo for pforHI/GF promoter | ACCCCCTGTGCACGAAGCCTGCGTGATTCATCGGCTGCGC |
| pforHI/GF F5 | SPR oligo for pforHI/GF promoter with linker | GCGCAGCCGATGAATCACGCAGGCTTCGTGCACAGGGGGTcctaccctacgtcctcctgc |
| pforHI/GF F6 | SPR oligo for pforHI/GF promoter | ATTCATCGGCTGCGCCCGACCCTAATCGCGGCCGGTGGGG |
| pforHI/GF F6 | SPR oligo for pforHI/GF promoter with linker | CCCCACCGGCCGCGATTAGGGTCGGGCGCAGCCGATGAATcctaccctacgtcctcctgc |
| pforHI/GF F7 | SPR oligo for pforHI/GF promoter | TCGCGGCCGGTGGGGGCGCTCAACGCATCCGGCCCAAGGA |
| pforHI/GF F7 | SPR oligo for pforHI/GF promoter with linker | TCCTTGGGCCGGATGCGTTGAGCGCCCCCACCGGCCGCGAcctaccctacgtcctcctgc |
| pforHI/GF F8 | SPR oligo for pforHI/GF promoter | CATCCGGCCCAAGGACAGATCCGGACAGTAGGGGGTGGCA |
| pforHI/GF F8 | SPR oligo for pforHI/GF promoter with linker | TGCCACCCCCTACTGTCCGGATCTGTCCTTGGGCCGGATGcctaccctacgtcctcctgc |
| pforHI/GF F9 | SPR oligo for pforHI/GF promoter | CAGTAGGGGGTGGCATGGCGGTCTATGGCACACGGGGGAG |
| pforHI/GF F9 | SPR oligo for pforHI/GF promoter with linker | CTCCCCCGTGTGCCATAGACCGCCATGCCACCCCCTACTGcctaccctacgtcctcctgc |
| pforHI/GF F10 | SPR oligo for pforHI/GF promoter | TGGCACACGGGGGAGGGGACATCCCTCCTTCGGCAGAGCC |
| pforHI/GF F10 | SPR oligo for pforHI/GF promoter with linker | GGCTCTGCCGAAGGAGGGATGTCCCCTCCCCCGTGTGCCAcctaccctacgtcctcctgc |
| pforHI/GF F11 | SPR oligo for pforHI/GF promoter | TCCTTCGGCAGAGCCGCCGCGGGTCGGACGGGGGACGGAA |
| pforHI/GF F11 | SPR oligo for pforHI/GF promoter with linker | TTCCGTCCCCCGTCCGACCCGCGGCGGCTCTGCCGAAGGAcctaccctacgtcctcctgc |
| pforHI/GF F12 | SPR oligo for pforHI/GF promoter | GGACGGGGGACGGAACCTGTCCCAGTGGGCGGGGCCGCTG |
| pforHI/GF F12 | SPR oligo for pforHI/GF promoter with linker | CAGCGGCCCCGCCCACTGGGACAGGTTCCGTCCCCCGTCCcctaccctacgtcctcctgc |
| pforHI/GF F13 | SPR oligo for pforHI/GF promoter | TGGGCGGGGCCGCTGTCCGAATCGGCATATTGCGACATTG |
| pforHI/GF F13 | SPR oligo for pforHI/GF promoter with linker | CAATGTCGCAATATGCCGATTCGGACAGCGGCCCCGCCCAcctaccctacgtcctcctgc |
| pforHI/GF F14 | SPR oligo for pforHI/GF promoter | CATATTGCGACATTGACGGCGTGCGGGATGACCGGAGTAA |
| pforHI/GF F14 | SPR oligo for pforHI/GF promoter with linker | TTACTCCGGTCATCCCGCACGCCGTCAATGTCGCAATATGcctaccctacgtcctcctgc |
| pforHI/GF F15 | SPR oligo for pforHI/GF promoter | GGATGACCGGAGTAACGTCACTCGTAGCGTGGCGGGACCC |
| pforHI/GF F15 | SPR oligo for pforHI/GF promoter with linker | GGGTCCCGCCACGCTACGAGTGACGTTACTCCGGTCATCCcctaccctacgtcctcctgc |
| pforHI/GF F16 | SPR oligo for pforHI/GF promoter | AGCGTGGCGGGACCCTCAGGTGACACTTACCTGCCTGCCG |
| pforHI/GF F16 | SPR oligo for pforHI/GF promoter with linker | CGGCAGGCAGGTAAGTGTCACCTGAGGGTCCCGCCACGCTcctaccctacgtcctcctgc |
| pforHI/GF F17 | SPR oligo for pforHI/GF promoter | CTTACCTGCCTGCCGCGGGCCGAAGGCGGAGGGGCAGTGC |
| pforHI/GF F17 | SPR oligo for pforHI/GF promoter with linker | GCACTGCCCCTCCGCCTTCGGCCCGCGGCAGGCAGGTAAGcctaccctacgtcctcctgc |
| pforHI/GF F18 | SPR oligo for pforHI/GF promoter | GCGGAGGGGCAGTGCTGAACGACGATGATCGCGCACTCGA |
| pforHI/GF F18 | SPR oligo for pforHI/GF promoter with linker | TCGAGTGCGCGATCATCGTCGTTCAGCACTGCCCCTCCGCcctaccctacgtcctcctgc |
| pforHI/GF F19 | SPR oligo for pforHI/GF promoter | TGATCGCGCACTCGACCAGTACGGGGGACCGTGACCGGCA |
| pforHI/GF F19 | SPR oligo for pforHI/GF promoter with linker | TGCCGGTCACGGTCCCCCGTACTGGTCGAGTGCGCGATCAcctaccctacgtcctcctgc |
| pforHI/GF F20 | SPR oligo for pforHI/GF promoter | GGACCGTGACCGGCACACAGTGGTGGCCTGCCCGTGGGCC |
| pforHI/GF F20 | SPR oligo for pforHI/GF promoter with linker | GGCCCACGGGCAGGCCACCACTGTGTGCCGGTCACGGTCCcctaccctacgtcctcctgc |
| pforHI/GF F21 | SPR oligo for pforHI/GF promoter | GCCTGCCCGTGGGCCCGCACCCGCCCGGCCGGCGGGAGTC |
| pforHI/GF F21 | SPR oligo for pforHI/GF promoter with linker | GACTCCCGCCGGCCGGGCGGGTGCGGGCCCACGGGCAGGCcctaccctacgtcctcctgc |
| pforHI/GF F22 | Oligo to narrow down ForF binding site | TGGCACACGGGGGAGGGGACATCCCTCCTTCGGCAGAGCC |
| pforHI/GF F22 | Oligo to narrow down ForF binding site with linker | GGCTCTGCCGAAGGAGGGATGTCCCCTCCCCCGTGTGCCAcctaccctacgtcctcctgc |
| pforHI/GF F23 | Oligo to narrow down ForF binding site | GCACACGGGGGAGGGGACATCCCTCCTTCGGCAGAGCC |
| pforHI/GF F23 | Oligo to narrow down ForF binding site with linker | GGCTCTGCCGAAGGAGGGATGTCCCCTCCCCCGTGTGCcctaccctacgtcctcctgc |
| pforHI/GF F24 | Oligo to narrow down ForF binding site | ACACGGGGGAGGGGACATCCCTCCTTCGGCAGAGCC |
| pforHI/GF F24 | Oligo to narrow down ForF binding site with linker | GGCTCTGCCGAAGGAGGGATGTCCCCTCCCCCGTGTcctaccctacgtcctcctgc |
| pforHI/GF F25 | Oligo to narrow down ForF binding site | ACGGGGGAGGGGACATCCCTCCTTCGGCAGAGCC |
| pforHI/GF F25 | Oligo to narrow down ForF binding site with linker | GGCTCTGCCGAAGGAGGGATGTCCCCTCCCCCGTcctaccctacgtcctcctgc |
| pforHI/GF F26 | Oligo to narrow down ForF binding site | GGGGGAGGGGACATCCCTCCTTCGGCAGAGCC |
| pforHI/GF F26 | Oligo to narrow down ForF binding site with linker | GGCTCTGCCGAAGGAGGGATGTCCCCTCCCCCcctaccctacgtcctcctgc |
| pforHI/GF F27 | Oligo to narrow down ForF binding site | GGGAGGGGACATCCCTCCTTCGGCAGAGCC |
| pforHI/GF F27 | Oligo to narrow down ForF binding site with linker | GGCTCTGCCGAAGGAGGGATGTCCCCTCCCcctaccctacgtcctcctgc |
| pforHI/GF F28 | Oligo to narrow down ForF binding site | GAGGGGACATCCCTCCTTCGGCAGAGCC |
| pforHI/GF F28 | Oligo to narrow down ForF binding site with linker | GGCTCTGCCGAAGGAGGGATGTCCCCTCcctaccctacgtcctcctgc |
| pforHI/GF F29 | Oligo to narrow down ForF binding site | GGGGACATCCCTCCTTCGGCAGAGCC |
| pforHI/GF F29 | Oligo to narrow down ForF binding site with linker | GGCTCTGCCGAAGGAGGGATGTCCCCcctaccctacgtcctcctgc |
| pforHI/GF F30 | Oligo to narrow down ForF binding site | GGACATCCCTCCTTCGGCAGAGCC |
| pforHI/GF F30 | Oligo to narrow down ForF binding site with linker | GGCTCTGCCGAAGGAGGGATGTCCcctaccctacgtcctcctgc |
| pforHI/GF F31 | Oligo to narrow down ForF binding site | ACATCCCTCCTTCGGCAGAGCC |
| pforHI/GF F31 | Oligo to narrow down ForF binding site with linker | GGCTCTGCCGAAGGAGGGATGTcctaccctacgtcctcctgc |
| pforHI/GF F32 | Oligo to narrow down ForF binding site | ATCCCTCCTTCGGCAGAGCC |
| pforHI/GF F32 | Oligo to narrow down ForF binding site with linker | GGCTCTGCCGAAGGAGGGATcctaccctacgtcctcctgc |
| pforHI/GF F33 | Oligo to narrow down ForF binding site | CCCTCCTTCGGCAGAGCC |
| pforHI/GF F33 | Oligo to narrow down ForF binding site with linker | GGCTCTGCCGAAGGAGGGcctaccctacgtcctcctgc |
| pforHI/GF F34 | Oligo to narrow down ForF binding site | CTCCTTCGGCAGAGCC |
| pforHI/GF F34 | Oligo to narrow down ForF binding site with linker | GGCTCTGCCGAAGGAGcctaccctacgtcctcctgc |
| pforHI/GF F35 | Oligo to narrow down ForF binding site | TGGCACACGGGGGAGG |
| pforHI/GF F35 | Oligo to narrow down ForF binding site with linker | CCTCCCCCGTGTGCCAcctaccctacgtcctcctgc |
| pforHI/GF F36 | Oligo to narrow down ForF binding site | TGGCACACGGGGGAGGGG |
| pforHI/GF F36 | Oligo to narrow down ForF binding site with linker | CCCCTCCCCCGTGTGCCAcctaccctacgtcctcctgc |
| pforHI/GF F37 | Oligo to narrow down ForF binding site | TGGCACACGGGGGAGGGGAC |
| pforHI/GF F37 | Oligo to narrow down ForF binding site with linker | GTCCCCTCCCCCGTGTGCCAcctaccctacgtcctcctgc |
| pforHI/GF F38 | Oligo to narrow down ForF binding site | TGGCACACGGGGGAGGGGACAT |
| pforHI/GF F38 | Oligo to narrow down ForF binding site with linker | ATGTCCCCTCCCCCGTGTGCCAcctaccctacgtcctcctgc |
| pforHI/GF F39 | Oligo to narrow down ForF binding site | TGGCACACGGGGGAGGGGACATCC |
| pforHI/GF F39 | Oligo to narrow down ForF binding site with linker | GGATGTCCCCTCCCCCGTGTGCCAcctaccctacgtcctcctgc |
| pforHI/GF F40 | Oligo to narrow down ForF binding site | TGGCACACGGGGGAGGGGACATCCCT |
| pforHI/GF F40 | Oligo to narrow down ForF binding site with linker | AGGGATGTCCCCTCCCCCGTGTGCCAcctaccctacgtcctcctgc |
| pforHI/GF F41 | Oligo to narrow down ForF binding site | TGGCACACGGGGGAGGGGACATCCCTCC |
| pforHI/GF F41 | Oligo to narrow down ForF binding site with linker | GGAGGGATGTCCCCTCCCCCGTGTGCCAcctaccctacgtcctcctgc |
| pforHI/GF F42 | Oligo to narrow down ForF binding site | TGGCACACGGGGGAGGGGACATCCCTCCTT |
| pforHI/GF F42 | Oligo to narrow down ForF binding site with linker | AAGGAGGGATGTCCCCTCCCCCGTGTGCCAcctaccctacgtcctcctgc |
| pforHI/GF F43 | Oligo to narrow down ForF binding site | TGGCACACGGGGGAGGGGACATCCCTCCTTCG |
| pforHI/GF F43 | Oligo to narrow down ForF binding site with linker | CGAAGGAGGGATGTCCCCTCCCCCGTGTGCCAcctaccctacgtcctcctgc |
| pforHI/GF F44 | Oligo to narrow down ForF binding site | TGGCACACGGGGGAGGGGACATCCCTCCTTCGGC |
| pforHI/GF F44 | Oligo to narrow down ForF binding site with linker | GCCGAAGGAGGGATGTCCCCTCCCCCGTGTGCCAcctaccctacgtcctcctgc |
| pforHI/GF F45 | Oligo to narrow down ForF binding site | TGGCACACGGGGGAGGGGACATCCCTCCTTCGGCAG |
| pforHI/GF F45 | Oligo to narrow down ForF binding site with linker | CTGCCGAAGGAGGGATGTCCCCTCCCCCGTGTGCCAcctaccctacgtcctcctgc |
| pforHI/GF F46 | Oligo to narrow down ForF binding site | TGGCACACGGGGGAGGGGACATCCCTCCTTCGGCAGAG |
| pforHI/GF F46 | Oligo to narrow down ForF binding site with linker | CTCTGCCGAAGGAGGGATGTCCCCTCCCCCGTGTGCCAcctaccctacgtcctcctgc |
| pKY5_0375 F1 | SPR oligo for pKY5_0375 promoter | GCGAAGGACCCGCCGCGTCACCGTGCCACCCCCGTCACAG |
| pKY5_0375 R1 | SPR oligo for pKY5_0375 promoter with linker | CTGTGACGGGGGTGGCACGGTGACGCGGCGGGTCCTTCGCcctaccctacgtcctcctgc |
| pKY5_0375 F2 | SPR oligo for pKY5_0375 promoter | CCACCCCCGTCACAGAGTTGATCACGCCCCTCGGCCAGAC |
| pKY5_0375 R2 | SPR oligo for pKY5_0375 promoter with linker | GTCTGGCCGAGGGGCGTGATCAACTCTGTGACGGGGGTGGcctaccctacgtcctcctgc |
| pKY5_0375 F3 | SPR oligo for pKY5_0375 promoter | GCCCCTCGGCCAGACCCCTGTGGCTATGCGCCCGTTGACC |
| pKY5_0375 R3 | SPR oligo for pKY5_0375 promoter with linker | GGTCAACGGGCGCATAGCCACAGGGGTCTGGCCGAGGGGCcctaccctacgtcctcctgc |
| pKY5_0375 F4 | SPR oligo for pKY5_0375 promoter | ATGCGCCCGTTGACCGCCACCGAAGGCCCGAAACCCGTCG |
| pKY5_0375 R4 | SPR oligo for pKY5_0375 promoter with linker | CGACGGGTTTCGGGCCTTCGGTGGCGGTCAACGGGCGCATcctaccctacgtcctcctgc |
| pKY5_0375 F5 | SPR oligo for pKY5_0375 promoter | GCCCGAAACCCGTCGCGCCGCACTCTCCGGAACGTCCGGA |
| pKY5_0375 R5 | SPR oligo for pKY5_0375 promoter with linker | TCCGGACGTTCCGGAGAGTGCGGCGCGACGGGTTTCGGGCcctaccctacgtcctcctgc |
| pKY5_0375 F6 | SPR oligo for pKY5_0375 promoter | TCCGGAACGTCCGGAGCCGTGTTGAAACGGACCAGACGAG |
| pKY5_0375 R6 | SPR oligo for pKY5_0375 promoter with linker | CTCGTCTGGTCCGTTTCAACACGGCTCCGGACGTTCCGGAcctaccctacgtcctcctgc |
| pKY5_0375 F7 | SPR oligo for pKY5_0375 promoter | AACGGACCAGACGAGCTACCCCGAACGGCGTAGCGAAGGG |
| pKY5_0375 R7 | SPR oligo for pKY5_0375 promoter with linker | CCCTTCGCTACGCCGTTCGGGGTAGCTCGTCTGGTCCGTTcctaccctacgtcctcctgc |
| pKY5_0375 F8 | SPR oligo for pKY5_0375 promoter | CGGCGTAGCGAAGGGTCCGCCAAGGCGTGGTCGAGAGTTG |
| pKY5_0375 R8 | SPR oligo for pKY5_0375 promoter with linker | CAACTCTCGACCACGCCTTGGCGGACCCTTCGCTACGCCGcctaccctacgtcctcctgc |
| pKY5_0375 F9 | SPR oligo for pKY5_0375 promoter | CGTGGTCGAGAGTTGCGTCAGAGGTGTACGAGGAGTGCGG |
| pKY5_0375 R9 | SPR oligo for pKY5_0375 promoter with linker | CCGCACTCCTCGTACACCTCTGACGCAACTCTCGACCACGcctaccctacgtcctcctgc |
| pKY5_0375 F10 | SPR oligo for pKY5_0375 promoter | GTACGAGGAGTGCGGAACGGACCATTCCGGTGCTGGTCGA |
| pKY5_0375 R10 | SPR oligo for pKY5_0375 promoter with linker | TCGACCAGCACCGGAATGGTCCGTTCCGCACTCCTCGTACcctaccctacgtcctcctgc |
| pKY5_0375 F11 | SPR oligo for pKY5_0375 promoter | TCCGGTGCTGGTCGAACCGTCGTCGGCCACCCGGACGGCT |
| pKY5_0375 R11 | SPR oligo for pKY5_0375 promoter with linker | AGCCGTCCGGGTGGCCGACGACGGTTCGACCAGCACCGGAcctaccctacgtcctcctgc |
| pKY5_0375 F12 | SPR oligo for pKY5_0375 promoter | GCCACCCGGACGGCTGCCCCTTTCGGGCGGTGTCCACCTG |
| pKY5_0375 R12 | SPR oligo for pKY5_0375 promoter with linker | CAGGTGGACACCGCCCGAAAGGGGCAGCCGTCCGGGTGGCcctaccctacgtcctcctgc |
| pKY5_0375 F13 | SPR oligo for pKY5_0375 promoter | GGCGGTGTCCACCTGAACGGGTCAGTCGTGAGGAGCCTCA |
| pKY5_0375 R13 | SPR oligo for pKY5_0375 promoter with linker | TGAGGCTCCTCACGACTGACCCGTTCAGGTGGACACCGCCcctaccctacgtcctcctgc |
| pKY5_0375 F14 | SPR oligo for pKY5_0375 promoter | TCGTGAGGAGCCTCACAAAACGCCGTTTCGGCTCGGTTTT |
| pKY5_0375 R14 | SPR oligo for pKY5_0375 promoter with linker | AAAACCGAGCCGAAACGGCGTTTTGTGAGGCTCCTCACGAcctaccctacgtcctcctgc |
| pKY5_0375 F15 | SPR oligo for pKY5_0375 promoter | TTTCGGCTCGGTTTTTTCGCCCTTGTCGGGTCAAGATCCT |
| pKY5_0375 R15 | SPR oligo for pKY5_0375 promoter with linker | AGGATCTTGACCCGACAAGGGCGAAAAAACCGAGCCGAAAcctaccctacgtcctcctgc |
| pKY5_0375 F16 | SPR oligo for pKY5_0375 promoter | TCGGGTCAAGATCCTCTCCGACGACAAGCCCCCGCCACAG |
| pKY5_0375 R16 | SPR oligo for pKY5_0375 promoter with linker | CTGTGGCGGGGGCTTGTCGTCGGAGAGGATCTTGACCCGAcctaccctacgtcctcctgc |
| pKY5_0375 F17 | Oligo to narrow down ForF binding site | AACGGACCAGACGAGCTACCCCGAACGGCGTAGCGAAGGG |
| pKY5_0375 R17 | Oligo to narrow down ForF binding site with linker | CCCTTCGCTACGCCGTTCGGGGTAGCTCGTCTGGTCCGTTcctaccctacgtcctcctgc |
| pKY5_0375 F18 | Oligo to narrow down ForF binding site | CGGACCAGACGAGCTACCCCGAACGGCGTAGCGAAGGG |
| pKY5_0375 R18 | Oligo to narrow down ForF binding site with linker | CCCTTCGCTACGCCGTTCGGGGTAGCTCGTCTGGTCCGcctaccctacgtcctcctgc |
| pKY5_0375 F19 | Oligo to narrow down ForF binding site | GACCAGACGAGCTACCCCGAACGGCGTAGCGAAGGG |
| pKY5_0375 R19 | Oligo to narrow down ForF binding site with linker | CCCTTCGCTACGCCGTTCGGGGTAGCTCGTCTGGTCcctaccctacgtcctcctgc |
| pKY5_0375 F20 | Oligo to narrow down ForF binding site | CCAGACGAGCTACCCCGAACGGCGTAGCGAAGGG |
| pKY5_0375 R20 | Oligo to narrow down ForF binding site with linker | CCCTTCGCTACGCCGTTCGGGGTAGCTCGTCTGGcctaccctacgtcctcctgc |
| pKY5_0375 F21 | Oligo to narrow down ForF binding site | AGACGAGCTACCCCGAACGGCGTAGCGAAGGG |
| pKY5_0375 R21 | Oligo to narrow down ForF binding site with linker | CCCTTCGCTACGCCGTTCGGGGTAGCTCGTCTcctaccctacgtcctcctgc |
| pKY5_0375 F22 | Oligo to narrow down ForF binding site | ACGAGCTACCCCGAACGGCGTAGCGAAGGG |
| pKY5_0375 R22 | Oligo to narrow down ForF binding site with linker | CCCTTCGCTACGCCGTTCGGGGTAGCTCGTcctaccctacgtcctcctgc |
| pKY5_0375 F23 | Oligo to narrow down ForF binding site | GAGCTACCCCGAACGGCGTAGCGAAGGG |
| pKY5_0375 R23 | Oligo to narrow down ForF binding site with linker | CCCTTCGCTACGCCGTTCGGGGTAGCTCcctaccctacgtcctcctgc |
| pKY5_0375 F24 | Oligo to narrow down ForF binding site | GCTACCCCGAACGGCGTAGCGAAGGG |
| pKY5_0375 R24 | Oligo to narrow down ForF binding site with linker | CCCTTCGCTACGCCGTTCGGGGTAGCcctaccctacgtcctcctgc |
| pKY5_0375 F25 | Oligo to narrow down ForF binding site | TACCCCGAACGGCGTAGCGAAGGG |
| pKY5_0375 R25 | Oligo to narrow down ForF binding site with linker | CCCTTCGCTACGCCGTTCGGGGTAcctaccctacgtcctcctgc |
| pKY5_0375 F26 | Oligo to narrow down ForF binding site | CCCCGAACGGCGTAGCGAAGGG |
| pKY5_0375 R26 | Oligo to narrow down ForF binding site with linker | CCCTTCGCTACGCCGTTCGGGGcctaccctacgtcctcctgc |
| pKY5_0375 F27 | Oligo to narrow down ForF binding site | CCGAACGGCGTAGCGAAGGG |
| pKY5_0375 R27 | Oligo to narrow down ForF binding site with linker | CCCTTCGCTACGCCGTTCGGcctaccctacgtcctcctgc |
| pKY5_0375 F28 | Oligo to narrow down ForF binding site | GAACGGCGTAGCGAAGGG |
| pKY5_0375 R28 | Oligo to narrow down ForF binding site with linker | CCCTTCGCTACGCCGTTCcctaccctacgtcctcctgc |
| pKY5_0375 F29 | Oligo to narrow down ForF binding site | ACGGCGTAGCGAAGGG |
| pKY5_0375 R29 | Oligo to narrow down ForF binding site with linker | CCCTTCGCTACGCCGTcctaccctacgtcctcctgc |
| pKY5_0375 F30 | Oligo to narrow down ForF binding site | AACGGACCAGACGAGC |
| pKY5_0375 R30 | Oligo to narrow down ForF binding site with linker | GCTCGTCTGGTCCGTTcctaccctacgtcctcctgc |
| pKY5_0375 F31 | Oligo to narrow down ForF binding site | AACGGACCAGACGAGCTA |
| pKY5_0375 R31 | Oligo to narrow down ForF binding site with linker | TAGCTCGTCTGGTCCGTTcctaccctacgtcctcctgc |
| pKY5_0375 F32 | Oligo to narrow down ForF binding site | AACGGACCAGACGAGCTACC |
| pKY5_0375 R32 | Oligo to narrow down ForF binding site with linker | GGTAGCTCGTCTGGTCCGTTcctaccctacgtcctcctgc |
| pKY5_0375 F33 | Oligo to narrow down ForF binding site | AACGGACCAGACGAGCTACCCC |
| pKY5_0375 R33 | Oligo to narrow down ForF binding site with linker | GGGGTAGCTCGTCTGGTCCGTTcctaccctacgtcctcctgc |
| pKY5_0375 F34 | Oligo to narrow down ForF binding site | AACGGACCAGACGAGCTACCCCGA |
| pKY5_0375 R34 | Oligo to narrow down ForF binding site with linker | TCGGGGTAGCTCGTCTGGTCCGTTcctaccctacgtcctcctgc |
| pKY5_0375 F35 | Oligo to narrow down ForF binding site | AACGGACCAGACGAGCTACCCCGAAC |
| pKY5_0375 R35 | Oligo to narrow down ForF binding site with linker | GTTCGGGGTAGCTCGTCTGGTCCGTTcctaccctacgtcctcctgc |
| pKY5_0375 F36 | Oligo to narrow down ForF binding site | AACGGACCAGACGAGCTACCCCGAACGG |
| pKY5_0375 R36 | Oligo to narrow down ForF binding site with linker | CCGTTCGGGGTAGCTCGTCTGGTCCGTTcctaccctacgtcctcctgc |
| pKY5_0375 F37 | Oligo to narrow down ForF binding site | AACGGACCAGACGAGCTACCCCGAACGGCG |
| pKY5_0375 R37 | Oligo to narrow down ForF binding site with linker | CGCCGTTCGGGGTAGCTCGTCTGGTCCGTTcctaccctacgtcctcctgc |
| pKY5_0375 F38 | Oligo to narrow down ForF binding site | AACGGACCAGACGAGCTACCCCGAACGGCGTA |
| pKY5_0375 R38 | Oligo to narrow down ForF binding site with linker | TACGCCGTTCGGGGTAGCTCGTCTGGTCCGTTcctaccctacgtcctcctgc |
| pKY5_0375 F39 | Oligo to narrow down ForF binding site | AACGGACCAGACGAGCTACCCCGAACGGCGTAGC |
| pKY5_0375 R39 | Oligo to narrow down ForF binding site with linker | GCTACGCCGTTCGGGGTAGCTCGTCTGGTCCGTTcctaccctacgtcctcctgc |
| pKY5_0375 F40 | Oligo to narrow down ForF binding site | AACGGACCAGACGAGCTACCCCGAACGGCGTAGCGA |
| pKY5_0375 R40 | Oligo to narrow down ForF binding site with linker | TCGCTACGCCGTTCGGGGTAGCTCGTCTGGTCCGTTcctaccctacgtcctcctgc |
| pKY5_0375 F41 | Oligo to narrow down ForF binding site | AACGGACCAGACGAGCTACCCCGAACGGCGTAGCGAAG |
| pKY5_0375 R41 | Oligo to narrow down ForF binding site with linker | CTTCGCTACGCCGTTCGGGGTAGCTCGTCTGGTCCGTTcctaccctacgtcctcctgc |
| pKY5_0375 F42 | Oligo to narrow down ForF binding site | GCCACCCGGACGGCTGCCCCTTTCGGGCGGTGTCCACCTG |
| pKY5_0375 R42 | Oligo to narrow down ForF binding site with linker | CAGGTGGACACCGCCCGAAAGGGGCAGCCGTCCGGGTGGCcctaccctacgtcctcctgc |
| pKY5_0375 F43 | Oligo to narrow down ForF binding site | CACCCGGACGGCTGCCCCTTTCGGGCGGTGTCCACCTG |
| pKY5_0375 R43 | Oligo to narrow down ForF binding site with linker | CAGGTGGACACCGCCCGAAAGGGGCAGCCGTCCGGGTGcctaccctacgtcctcctgc |
| pKY5_0375 F44 | Oligo to narrow down ForF binding site | CCCGGACGGCTGCCCCTTTCGGGCGGTGTCCACCTG |
| pKY5_0375 R44 | Oligo to narrow down ForF binding site with linker | CAGGTGGACACCGCCCGAAAGGGGCAGCCGTCCGGGcctaccctacgtcctcctgc |
| pKY5_0375 F45 | Oligo to narrow down ForF binding site | CGGACGGCTGCCCCTTTCGGGCGGTGTCCACCTG |
| pKY5_0375 R45 | Oligo to narrow down ForF binding site with linker | CAGGTGGACACCGCCCGAAAGGGGCAGCCGTCCGcctaccctacgtcctcctgc |
| pKY5_0375 F46 | Oligo to narrow down ForF binding site | GACGGCTGCCCCTTTCGGGCGGTGTCCACCTG |
| pKY5_0375 R46 | Oligo to narrow down ForF binding site with linker | CAGGTGGACACCGCCCGAAAGGGGCAGCCGTCcctaccctacgtcctcctgc |
| pKY5_0375 F47 | Oligo to narrow down ForF binding site | CGGCTGCCCCTTTCGGGCGGTGTCCACCTG |
| pKY5_0375 R47 | Oligo to narrow down ForF binding site with linker | CAGGTGGACACCGCCCGAAAGGGGCAGCCGcctaccctacgtcctcctgc |
| pKY5_0375 F48 | Oligo to narrow down ForF binding site | GCTGCCCCTTTCGGGCGGTGTCCACCTG |
| pKY5_0375 R48 | Oligo to narrow down ForF binding site with linker | CAGGTGGACACCGCCCGAAAGGGGCAGCcctaccctacgtcctcctgc |
| pKY5_0375 F49 | Oligo to narrow down ForF binding site | TGCCCCTTTCGGGCGGTGTCCACCTG |
| pKY5_0375 R49 | Oligo to narrow down ForF binding site with linker | CAGGTGGACACCGCCCGAAAGGGGCAcctaccctacgtcctcctgc |
| pKY5_0375 F50 | Oligo to narrow down ForF binding site | CCCCTTTCGGGCGGTGTCCACCTG |
| pKY5_0375 R50 | Oligo to narrow down ForF binding site with linker | CAGGTGGACACCGCCCGAAAGGGGcctaccctacgtcctcctgc |
| pKY5_0375 F51 | Oligo to narrow down ForF binding site | CCTTTCGGGCGGTGTCCACCTG |
| pKY5_0375 R51 | Oligo to narrow down ForF binding site with linker | CAGGTGGACACCGCCCGAAAGGcctaccctacgtcctcctgc |
| pKY5_0375 F52 | Oligo to narrow down ForF binding site | TTTCGGGCGGTGTCCACCTG |
| pKY5_0375 R52 | Oligo to narrow down ForF binding site with linker | CAGGTGGACACCGCCCGAAAcctaccctacgtcctcctgc |
| pKY5_0375 F53 | Oligo to narrow down ForF binding site | TCGGGCGGTGTCCACCTG |
| pKY5_0375 R53 | Oligo to narrow down ForF binding site with linker | CAGGTGGACACCGCCCGAcctaccctacgtcctcctgc |
| pKY5_0375 F54 | Oligo to narrow down ForF binding site | GGGCGGTGTCCACCTG |
| pKY5_0375 R54 | Oligo to narrow down ForF binding site with linker | CAGGTGGACACCGCCCcctaccctacgtcctcctgc |
| pKY5_0375 F55 | Oligo to narrow down ForF binding site | GCCACCCGGACGGCTG |
| pKY5_0375 R55 | Oligo to narrow down ForF binding site with linker | CAGCCGTCCGGGTGGCcctaccctacgtcctcctgc |
| pKY5_0375 F56 | Oligo to narrow down ForF binding site | GCCACCCGGACGGCTGCC |
| pKY5_0375 R56 | Oligo to narrow down ForF binding site with linker | GGCAGCCGTCCGGGTGGCcctaccctacgtcctcctgc |
| pKY5_0375 F57 | Oligo to narrow down ForF binding site | GCCACCCGGACGGCTGCCCC |
| pKY5_0375 R57 | Oligo to narrow down ForF binding site with linker | GGGGCAGCCGTCCGGGTGGCcctaccctacgtcctcctgc |
| pKY5_0375 F58 | Oligo to narrow down ForF binding site | GCCACCCGGACGGCTGCCCCTT |
| pKY5_0375 R58 | Oligo to narrow down ForF binding site with linker | AAGGGGCAGCCGTCCGGGTGGCcctaccctacgtcctcctgc |
| pKY5_0375 F59 | Oligo to narrow down ForF binding site | GCCACCCGGACGGCTGCCCCTTTC |
| pKY5_0375 R59 | Oligo to narrow down ForF binding site with linker | GAAAGGGGCAGCCGTCCGGGTGGCcctaccctacgtcctcctgc |
| pKY5_0375 F60 | Oligo to narrow down ForF binding site | GCCACCCGGACGGCTGCCCCTTTCGG |
| pKY5_0375 R60 | Oligo to narrow down ForF binding site with linker | CCGAAAGGGGCAGCCGTCCGGGTGGCcctaccctacgtcctcctgc |
| pKY5_0375 F61 | Oligo to narrow down ForF binding site | GCCACCCGGACGGCTGCCCCTTTCGGGC |
| pKY5_0375 R61 | Oligo to narrow down ForF binding site with linker | GCCCGAAAGGGGCAGCCGTCCGGGTGGCcctaccctacgtcctcctgc |
| pKY5_0375 F62 | Oligo to narrow down ForF binding site | GCCACCCGGACGGCTGCCCCTTTCGGGCGG |
| pKY5_0375 R62 | Oligo to narrow down ForF binding site with linker | CCGCCCGAAAGGGGCAGCCGTCCGGGTGGCcctaccctacgtcctcctgc |
| pKY5_0375 F63 | Oligo to narrow down ForF binding site | GCCACCCGGACGGCTGCCCCTTTCGGGCGGTG |
| pKY5_0375 R63 | Oligo to narrow down ForF binding site with linker | CACCGCCCGAAAGGGGCAGCCGTCCGGGTGGCcctaccctacgtcctcctgc |
| pKY5_0375 F64 | Oligo to narrow down ForF binding site | GCCACCCGGACGGCTGCCCCTTTCGGGCGGTGTC |
| pKY5_0375 R64 | Oligo to narrow down ForF binding site with linker | GACACCGCCCGAAAGGGGCAGCCGTCCGGGTGGCcctaccctacgtcctcctgc |
| pKY5_0375 F65 | Oligo to narrow down ForF binding site | GCCACCCGGACGGCTGCCCCTTTCGGGCGGTGTCCA |
| pKY5_0375 R65 | Oligo to narrow down ForF binding site with linker | TGGACACCGCCCGAAAGGGGCAGCCGTCCGGGTGGCcctaccctacgtcctcctgc |
| pKY5_0375 F66 | Oligo to narrow down ForF binding site | GCCACCCGGACGGCTGCCCCTTTCGGGCGGTGTCCACC |
| pKY5_0375 R66 | Oligo to narrow down ForF binding site with linker | GGTGGACACCGCCCGAAAGGGGCAGCCGTCCGGGTGGCcctaccctacgtcctcctgc |
| pforZ/AA F1 | SPR oligo for pforZ/AA promoter | CTCGCCGACGACGTGAAGGCCCTCATCCGGCTCGTCACGC |
| pforZ/AA R1 | SPR oligo for pforZ/AA promoter with linker | GCGTGACGAGCCGGATGAGGGCCTTCACGTCGTCGGCGAGcctaccctacgtcctcctgc |
| pforZ/AA F2 | SPR oligo for pforZ/AA promoter | AGTCCCGAGCTCGCCGACGACGTGAAGGCCCTCATCCGGC |
| pforZ/AA R2 | SPR oligo for pforZ/AA promoter with linker | GCCGGATGAGGGCCTTCACGTCGTCGGCGAGCTCGGGACTcctaccctacgtcctcctgc |
| pforZ/AA F3 | SPR oligo for pforZ/AA promoter | CGCAGTCGAGTTGACCGCCGACGCGAGTCCCGAGCTCGCC |
| pforZ/AA R3 | SPR oligo for pforZ/AA promoter with linker | GGCGAGCTCGGGACTCGCGTCGGCGGTCAACTCGACTGCGcctaccctacgtcctcctgc |
| pforZ/AA F4 | SPR oligo for pforZ/AA promoter | CCGCCGGTGCCGAACCGGACGCAGCCGCAGTCGAGTTGAC |
| pforZ/AA R4 | SPR oligo for pforZ/AA promoter with linker | GTCAACTCGACTGCGGCTGCGTCCGGTTCGGCACCGGCGGcctaccctacgtcctcctgc |
| pforZ/AA F5 | SPR oligo for pforZ/AA promoter | GGACTTCAAGCCATGGCCTCCCCTCCCGCCGGTGCCGAAC |
| pforZ/AA R5 | SPR oligo for pforZ/AA promoter with linker | GTTCGGCACCGGCGGGAGGGGAGGCCATGGCTTGAAGTCCcctaccctacgtcctcctgc |
| pforZ/AA F6 | SPR oligo for pforZ/AA promoter | CTTGCAGTACAGCACCGGACGTGCTGGACTTCAAGCCATG |
| pforZ/AA R6 | SPR oligo for pforZ/AA promoter with linker | CATGGCTTGAAGTCCAGCACGTCCGGTGCTGTACTGCAAGcctaccctacgtcctcctgc |
| pforZ/AA F7 | SPR oligo for pforZ/AA promoter | CACATTAGCTCGAAGTTCGATGCATCTTGCAGTACAGCAC |
| pforZ/AA R7 | SPR oligo for pforZ/AA promoter with linker | GTGCTGTACTGCAAGATGCATCGAACTTCGAGCTAATGTGcctaccctacgtcctcctgc |
| pforZ/AA F8 | SPR oligo for pforZ/AA promoter | ACGGCCTCCCGGATTGCCTCTGAGTCACATTAGCTCGAAG |
| pforZ/AA R8 | SPR oligo for pforZ/AA promoter with linker | CTTCGAGCTAATGTGACTCAGAGGCAATCCGGGAGGCCGTcctaccctacgtcctcctgc |
| pforZ/AA F9 | SPR oligo for pforZ/AA promoter | CTCGTTCACGCCTTGGGCACCCATGACGGCCTCCCGGATT |
| pforZ/AA R9 | SPR oligo for pforZ/AA promoter with linker | AATCCGGGAGGCCGTCATGGGTGCCCAAGGCGTGAACGAGcctaccctacgtcctcctgc |
| pforZ/AA F10 | SPR oligo for pforZ/AA promoter | CCGGCGGGCCCTCGCGCTCCGGTAACTCGTTCACGCCTTG |
| pforZ/AA R10 | SPR oligo for pforZ/AA promoter with linker | CAAGGCGTGAACGAGTTACCGGAGCGCGAGGGCCCGCCGGcctaccctacgtcctcctgc |
| pforZ/AA F11 | SPR oligo for pforZ/AA promoter | TCCCTCCCCAGGCGTTCCTCCGGTCCCGGCGGGCCCTCGC |
| pforZ/AA R11 | SPR oligo for pforZ/AA promoter with linker | GCGAGGGCCCGCCGGGACCGGAGGAACGCCTGGGGAGGGAcctaccctacgtcctcctgc |
| pforZ/AA F12 | SPR oligo for pforZ/AA promoter | GACCACGCCGGCCGTCACCCAGATCTCCCTCCCCAGGCGT |
| pforZ/AA R12 | SPR oligo for pforZ/AA promoter with linker | ACGCCTGGGGAGGGAGATCTGGGTGACGGCCGGCGTGGTCcctaccctacgtcctcctgc |
| pforZ/AA F13 | SPR oligo for pforZ/AA promoter | GCACCGCCATGATCGTGCCGAGGACGACCACGCCGGCCGT |
| pforZ/AA R13 | SPR oligo for pforZ/AA promoter with linker | ACGGCCGGCGTGGTCGTCCTCGGCACGATCATGGCGGTGCcctaccctacgtcctcctgc |
| pforZ/AA F14 | SPR oligo for pforZ/AA promoter | GCGACGTTGACGATGGTGGTGTCGAGCACCGCCATGATCG |
| pforZ/AA R14 | SPR oligo for pforZ/AA promoter with linker | CGATCATGGCGGTGCTCGACACCACCATCGTCAACGTCGCcctaccctacgtcctcctgc |
| pforZ/AA F15 | SPR oligo for pforZ/AA promoter | GAGGTCCTTGCTCAGGGTGTTGAGGGCGACGTTGACGATG |
| pforZ/AA R15 | SPR oligo for pforZ/AA promoter with linker | CATCGTCAACGTCGCCCTCAACACCCTGAGCAAGGACCTCcctaccctacgtcctcctgc |
| pforZ/AA F16 | SPR oligo for pforZ/AA promoter | ATTGGAGCGTGGCGAGTCCGGTGTCGAGGTCCTTGCTCAG |
| pforZ/AA R16 | SPR oligo for pforZ/AA promoter with linker | CTGAGCAAGGACCTCGACACCGGACTCGCCACGCTCCAATcctaccctacgtcctcctgc |
| pforZ/AA F17 | SPR oligo for pforZ/AA promoter | GCGAGGAAGTAGCCGGTCACCACCCATTGGAGCGTGGCGA |
| pforZ/AA R17 | SPR oligo for pforZ/AA promoter with linker | TCGCCACGCTCCAATGGGTGGTGACCGGCTACTTCCTCGCcctaccctacgtcctcctgc |
| pforZ/AA F18 | Oligo to narrow down ForZ binding site | CCGGATTGCCTCTGAGTCAC |
| pforZ/AA R18 | Oligo to narrow down ForZ binding site with linker | GTGACTCAGAGGCAATCCGGcctaccctacgtcctcctgc |
| pforZ/AA F19 | Oligo to narrow down ForZ binding site | TTGCCTCTGAGTCACATTAG |
| pforZ/AA R19 | Oligo to narrow down ForZ binding site with linker | CTAATGTGACTCAGAGGCAAcctaccctacgtcctcctgc |
| pforZ/AA F20 | Oligo to narrow down ForZ binding site | TGAGTCACATTAGCTCGA |
| pforZ/AA R20 | Oligo to narrow down ForZ binding site with linker | TCGAGCTAATGTGACTCAcctaccctacgtcctcctgc |
| pforZ/AA F21 | Oligo to narrow down ForZ binding site | GTCACATTAGCTCGAAGTTC |
| pforZ/AA R21 | Oligo to narrow down ForZ binding site with linker | GAACTTCGAGCTAATGTGACcctaccctacgtcctcctgc |
| pforZ/AA F22 | Oligo to narrow down ForZ binding site | ATTAGCTCGAAGTTCGATGC |
| pforZ/AA R22 | Oligo to narrow down ForZ binding site with linker | GCATCGAACTTCGAGCTAATcctaccctacgtcctcctgc |
| pforZ/AA F23 | Oligo to narrow down ForZ binding site | TCGAAGTTCGATGCATCTT |
| pforZ/AA R23 | Oligo to narrow down ForZ binding site with linker | AAGATGCATCGAACTTCGAcctaccctacgtcctcctgc |
| pforZ/AA F24 | Oligo to narrow down ForZ binding site | AGTTCGATGCATCTTGCAGT |
| pforZ/AA R24 | Oligo to narrow down ForZ binding site with linker | ACTGCAAGATGCATCGAACTcctaccctacgtcctcctgc |
| pforZ/AA F25 | Oligo to narrow down ForZ binding site | GATGCATCTTGCAGTACAGC |
| pforZ/AA R25 | Oligo to narrow down ForZ binding site with linker | GCTGTACTGCAAGATGCATCcctaccctacgtcctcctgc |
| pforZ/AA F26 | Oligo to narrow down ForZ binding site | ATCTTGCAGTACAGCACCGG |
| pforZ/AA R26 | Oligo to narrow down ForZ binding site with linker | CCGGTGCTGTACTGCAAGATcctaccctacgtcctcctgc |
| pforZ/AA F27 | Oligo to narrow down ForZ binding site | CCGGATTGCCTCTGAGTCACATTAGCTCGA |
| pforZ/AA R27 | Oligo to narrow down ForZ binding site with linker | CCGGATTGCCTCTGAGTCACATTAGCTCGA |
| pforZ/AA F28 | Oligo to narrow down ForZ binding site | TCGAGCTAATGTGACTCAGAGGCAATCCGGcctaccctacgtcctcctgc |
| pforZ/AA R28 | Oligo to narrow down ForZ binding site with linker | TTGCCTCTGAGTCACATTAGCTCGAAGTTC |
| pforZ/AA F29 | Oligo to narrow down ForZ binding site | GAACTTCGAGCTAATGTGACTCAGAGGCAAcctaccctacgtcctcctgc |
| pforZ/AA R29 | Oligo to narrow down ForZ binding site with linker | TCTGAGTCACATTAGCTCGAAGTTCGATGC |
| pforZ/AA F30 | Oligo to narrow down ForZ binding site | GCATCGAACTTCGAGCTAATGTGACTCAGAcctaccctacgtcctcctgc |
| pforZ/AA R30 | Oligo to narrow down ForZ binding site with linker | GTCACATTAGCTCGAAGTTCGATGCATCTT |
| pforZ/AA F31 | Oligo to narrow down ForZ binding site | AAGATGCATCGAACTTCGAGCTAATGTGACcctaccctacgtcctcctgc |
| pforZ/AA R31 | Oligo to narrow down ForZ binding site with linker | ATTAGCTCGAAGTTCGATGCATCTTGCAGT |
| pforZ/AA F32 | Oligo to narrow down ForZ binding site | ACTGCAAGATGCATCGAACTTCGAGCTAATcctaccctacgtcctcctgc |
| pforZ/AA R32 | Oligo to narrow down ForZ binding site with linker | CTCGAAGTTCGATGCATCTTGCAGTACAGC |
| pforZ/AA F33 | Oligo to narrow down ForZ binding site | GCTGTACTGCAAGATGCATCGAACTTCGAGcctaccctacgtcctcctgc |
| pforZ/AA R33 | Oligo to narrow down ForZ binding site with linker | AGTTCGATGCATCTTGCAGTACAGCACCGG |
| pforZ/AA F34 | Oligo to narrow down ForZ binding site | CCGGTGCTGTACTGCAAGATGCATCGAACTcctaccctacgtcctcctgc |
| pforZ/AA R34 | Oligo to narrow down ForZ binding site with linker | GATGCATCTTGCAGTACAGCACCGGACGTG |
| pforT/U F1 | SPR oligo for pforT/U promoter | CGAGGAGCGGGAAGGACGCCGCTTCGCGCACACCGACCTC |
| pforT/U R1 | SPR oligo for pforT/U promoter with linker | GAGGTCGGTGTGCGCGAAGCGGCGTCCTTCCCGCTCCTCGcctaccctacgtcctcctgc |
| pforT/U F2 | SPR oligo for pforT/U promoter | CACGCACGGCGTCTTCGAGGAGCGGGAAGGACGCCGCTTC |
| pforT/U R2 | SPR oligo for pforT/U promoter with linker | GAAGCGGCGTCCTTCCCGCTCCTCGAAGACGCCGTGCGTGcctaccctacgtcctcctgc |
| pforT/U F3 | SPR oligo for pforT/U promoter | GAAATCGACGCTCACGCACCGACGTTGGAGCGCCTGCTGC |
| pforT/U R3 | SPR oligo for pforT/U promoter with linker | GCAGCAGGCGCTCCAACGTCGGTGCGTGAGCGTCGATTTCcctaccctacgtcctcctgc |
| pforT/U F4 | SPR oligo for pforT/U promoter | GAAATCGACGCTCACGCACCGACGTTGGAGCGCCTGCTGC |
| pforT/U R4 | SPR oligo for pforT/U promoter with linker | GCAGCAGGCGCTCCAACGTCGGTGCGTGAGCGTCGATTTCcctaccctacgtcctcctgc |
| pforT/U F5 | SPR oligo for pforT/U promoter | GCGCACCGCGGCCGAGCTCGCCCCCGAAATCGACGCTCAC |
| pforT/U R5 | SPR oligo for pforT/U promoter with linker | GTGAGCGTCGATTTCGGGGGCGAGCTCGGCCGCGGTGCGCcctaccctacgtcctcctgc |
| pforT/U F6 | SPR oligo for pforT/U promoter | TCCCGGACGCGCTCGACGAGGAACCGCGCACCGCGGCCGA |
| pforT/U R6 | SPR oligo for pforT/U promoter with linker | TCGGCCGCGGTGCGCGGTTCCTCGTCGAGCGCGTCCGGGAcctaccctacgtcctcctgc |
| pforT/U F7 | SPR oligo for pforT/U promoter | CTGCGGGCCGCCGTCACGCTCGGCGTCCCGGACGCGCTCG |
| pforT/U R7 | SPR oligo for pforT/U promoter with linker | CGAGCGCGTCCGGGACGCCGAGCGTGACGGCGGCCCGCAGcctaccctacgtcctcctgc |
| pforT/U F8 | SPR oligo for pforT/U promoter | CATGAGCCTCGGCTTCGCCGGCGCCCTGCGGGCCGCCGTC |
| pforT/U R8 | SPR oligo for pforT/U promoter with linker | GACGGCGGCCCGCAGGGCGCCGGCGAAGCCGAGGCTCATGcctaccctacgtcctcctgc |
| pforT/U F9 | SPR oligo for pforT/U promoter | CCGCCACGAAACTCCGCGAGCTGGGCATGAGCCTCGGCTT |
| pforT/U R9 | SPR oligo for pforT/U promoter with linker | AAGCCGAGGCTCATGCCCAGCTCGCGGAGTTTCGTGGCGGcctaccctacgtcctcctgc |
| pforT/U F10 | SPR oligo for pforT/U promoter | ACTCCCTTGCGCCCCACGCCCGAAGCCGCCACGAAACTCC |
| pforT/U R10 | SPR oligo for pforT/U promoter with linker | GGAGTTTCGTGGCGGCTTCGGGCGTGGGGCGCAAGGGAGTcctaccctacgtcctcctgc |
| pforT/U F11 | SPR oligo for pforT/U promoter | GGAGGAGCACCTCACCATGACGTCAACTCCCTTGCGCCCC |
| pforT/U R11 | SPR oligo for pforT/U promoter with linker | GGGGCGCAAGGGAGTTGACGTCATGGTGAGGTGCTCCTCCcctaccctacgtcctcctgc |
| pforT/U F12 | SPR oligo for pforT/U promoter | GGATTGGTCCCCAAATTGCTTTACAGGAGGAGCACCTCAC |
| pforT/U R12 | SPR oligo for pforT/U promoter with linker | GTGAGGTGCTCCTCCTGTAAAGCAATTTGGGGACCAATCCcctaccctacgtcctcctgc |
| pforT/U F13 | SPR oligo for pforT/U promoter | AGAGTCGTTGATTGAGTAAGTCAATGGATTGGTCCCCAAA |
| pforT/U R13 | SPR oligo for pforT/U promoter with linker | TTTGGGGACCAATCCATTGACTTACTCAATCAACGACTCTcctaccctacgtcctcctgc |
| pforT/U F14 | SPR oligo for pforT/U promoter | TCTTGCGCGGCTTGAGCCATTATGTAGAGTCGTTGATTGA |
| pforT/U R14 | SPR oligo for pforT/U promoter with linker | TCAATCAACGACTCTACATAATGGCTCAAGCCGCGCAAGAcctaccctacgtcctcctgc |
| pforT/U F15 | SPR oligo for pforT/U promoter | AGCTGCCCTCACTCTCTCGGTTGCCTCTTGCGCGGCTTGA |
| pforT/U R15 | SPR oligo for pforT/U promoter with linker | TCAAGCCGCGCAAGAGGCAACCGAGAGAGTGAGGGCAGCTcctaccctacgtcctcctgc |
| pforT/U F16 | SPR oligo for pforT/U promoter | GTCGGCGGAGATCTCGGCCATGTGGAGCTGCCCTCACTCT |
| pforT/U R16 | SPR oligo for pforT/U promoter with linker | AGAGTGAGGGCAGCTCCACATGGCCGAGATCTCCGCCGACcctaccctacgtcctcctgc |
| pforT/U F17 | SPR oligo for pforT/U promoter | CACGTCCGGTGCTGTACTGCAAGATGCATCcctaccctacgtcctcctgc |
| pforT/U R17 | SPR oligo for pforT/U promoter with linker | GAGATCTCCGCCGACAACGAGTTCGCCACGTTCATCAACCcctaccctacgtcctcctgc |
| pforT/U F18 | Oligo to narrow down ForJ binding site | gccattatgtagagtcgttgattgagtaag |
| pforT/U R18 | Oligo to narrow down ForJ binding site with linker | cttactcaatcaacgactctacataatggCcctaccctacgtcctcctgc |
| pforT/U F19 | Oligo to narrow down ForJ binding site | Tatgtagagtcgttgattgagtaagtcaat |
| pforT/U R19 | Oligo to narrow down ForJ binding site with linker | attgacttactcaatcaacgactctacatAcctaccctacgtcctcctgc |
| pforT/U F20 | Oligo to narrow down ForJ binding site | Agagtcgttgattgagtaagtcaatggatt |
| pforT/U R20 | Oligo to narrow down ForJ binding site with linker | aatccattgacttactcaatcaacgactcTcctaccctacgtcctcctgc |
| pforT/U F21 | Oligo to narrow down ForJ binding site | Cgttgattgagtaagtcaatggattggtcc |
| pforT/U R21 | Oligo to narrow down ForJ binding site with linker | ggaccaatccattgacttactcaatcaacGcctaccctacgtcctcctgc |
| pforT/U F22 | Oligo to narrow down ForJ binding site | Attgagtaagtcaatggattggtccccaaa |
| pforT/U R22 | Oligo to narrow down ForJ binding site with linker | tttggggaccaatccattgacttactcaaTcctaccctacgtcctcctgc |
| pforT/U F23 | Oligo to narrow down ForJ binding site | Gtaagtcaatggattggtccccaaattgct |
| pforT/U R23 | Oligo to narrow down ForJ binding site with linker | agcaatttggggaccaatccattgacttaCcctaccctacgtcctcctgc |
| pforT/U F24 | Oligo to narrow down ForJ binding site | Tcaatggattggtccccaaattgctttaca |
| pforT/U R24 | Oligo to narrow down ForJ binding site with linker | tgtaaagcaatttggggaccaatccattgAcctaccctacgtcctcctgc |
| pforM F1 | SPR oligo for pforM promoter | ACACCGGCTCCCATCGGTTGCTGTGCTAATTGATTGAGTA |
| pforM R1 | SPR oligo for pforM promoter with linker | TACTCAATCAATTAGCACAGCAACCGATGGGAGCCGGTGTcctaccctacgtcctcctgc |
| pforM F2 | SPR oligo for pforM promoter | CTAATTGATTGAGTAATTGCTTGAGTTACTCTACTAAATA |
| pforM R2 | SPR oligo for pforM promoter with linker | TATTTAGTAGAGTAACTCAAGCAATTACTCAATCAATTAGcctaccctacgtcctcctgc |
| pforM F3 | SPR oligo for pforM promoter | TTACTCTACTAAATACCCGGATCCACGCAAGGGCCCGGGC |
| pforM R3 | SPR oligo for pforM promoter with linker | GCCCGGGCCCTTGCGTGGATCCGGGTATTTAGTAGAGTAAcctaccctacgtcctcctgc |
| pforM F4 | SPR oligo for pforM promoter | CGCAAGGGCCCGGGCCCACCCACGCAAGGGCCCGGGCCCG |
| pforM R4 | SPR oligo for pforM promoter with linker | CGGGCCCGGGCCCTTGCGTGGGTGGGCCCGGGCCCTTGCGcctaccctacgtcctcctgc |
| pforM F5 | SPR oligo for pforM promoter | AAGGGCCCGGGCCCGTCCACACAGGGGCCCGGGCCCGCGC |
| pforM R5 | SPR oligo for pforM promoter with linker | GCGCGGGCCCGGGCCCCTGTGTGGACGGGCCCGGGCCCTTcctaccctacgtcctcctgc |
| pforM F6 | SPR oligo for pforM promoter | GGCCCGGGCCCGCGCCCCGTCGCCGCGCCGAGTTCGCGTC |
| pforM R6 | SPR oligo for pforM promoter with linker | GACGCGAACTCGGCGCGGCGACGGGGCGCGGGCCCGGGCCcctaccctacgtcctcctgc |
| pforM F7 | SPR oligo for pforM promoter | CGCCGAGTTCGCGTCATGTTCACGCGCGGCTCATTGACGC |
| pforM R7 | SPR oligo for pforM promoter with linker | GCGTCAATGAGCCGCGCGTGAACATGACGCGAACTCGGCGcctaccctacgtcctcctgc |
| pforM F8 | SPR oligo for pforM promoter | GCGGCTCATTGACGCGCCTCGGCCACCTCCTTAGCTTCAT |
| pforM R8 | SPR oligo for pforM promoter with linker | ATGAAGCTAAGGAGGTGGCCGAGGCGCGTCAATGAGCCGCcctaccctacgtcctcctgc |
| pforM F9 | SPR oligo for pforM promoter | CCTCCTTAGCTTCATGGCGCATCTGTCATCACGCGGGAGG |
| pforM R9 | SPR oligo for pforM promoter with linker | CCTCCCGCGTGATGACAGATGCGCCATGAAGCTAAGGAGGcctaccctacgtcctcctgc |
| pforM F10 | SPR oligo for pforM promoter | TCATCACGCGGGAGGGGTCTGTCGT |
| pforM R10 | SPR oligo for pforM promoter with linker | ACGACAGACCCCTCCCGCGTGATGAcctaccctacgtcctcctgc |
| pforJ F1 | SPR oligo for pforJ promoter | GGCGCCGTGGTCGTGGTCATGTTCGCTCACCTCTGCTGTG |
| pforJ R1 | SPR oligo for pforJ promoter with linker | CACAGCAGAGGTGAGCGAACATGACCACGACCACGGCGCCcctaccctacgtcctcctgc |
| pforJ F2 | SPR oligo for pforJ promoter | CTCACCTCTGCTGTGACGTGCTTCGAGACCGCCGGTGGGG |
| pforJ R2 | SPR oligo for pforJ promoter with linker | CCCCACCGGCGGTCTCGAAGCACGTCACAGCAGAGGTGAGcctaccctacgtcctcctgc |
| pforJ F3 | SPR oligo for pforJ promoter | AGACCGCCGGTGGGGGAATCGGCGGACTTGGTCAATCATT |
| pforJ R3 | SPR oligo for pforJ promoter with linker | AATGATTGACCAAGTCCGCCGATTCCCCCACCGGCGGTCTcctaccctacgtcctcctgc |
| pforJ F4 | SPR oligo for pforJ promoter | ACTTGGTCAATCATTGCAGAAGTCAGTGGTGGATGGATAG |
| pforJ R4 | SPR oligo for pforJ promoter with linker | CTATCCATCCACCACTGACTTCTGCAATGATTGACCAAGTcctaccctacgtcctcctgc |
| pforJ F5 | SPR oligo for pforJ promoter | GTGGTGGATGGATAGAAGTG |
| pforJ R5 | SPR oligo for pforJ promoter with linker | CACTTCTATCCATCCACCACcctaccctacgtcctcctgc |
| forE F1 | SPR oligo for forE | CGCAGCCGGACGCCCCGCCCTTCGTGAAGCCGGGGGACGC |
| forE R1 | SPR oligo for forE with linker | GCGTCCCCCGGCTTCACGAAGGGCGGGGCGTCCGGCTGCGcctaccctacgtcctcctgc |
| forE F2 | SPR oligo for forE | GAAGCCGGGGGACGCCGTCACCCCCGGCCAGCAGATCGGC |
| forE R2 | SPR oligo for forE with linker | GCCGATCTGCTGGCCGGGGGTGACGGCGTCCCCCGGCTTCcctaccctacgtcctcctgc |
| forE F3 | SPR oligo for forE | GGCCAGCAGATCGGCGTCGTCGAGGCGATGAAGTTGATGA |
| forE R3 | SPR oligo for forE with linker | TCATCAACTTCATCGCCTCGACGACGCCGATCTGCTGGCCcctaccctacgtcctcctgc |
| forE F4 | SPR oligo for forE | CGATGAAGTTGATGACGCCGGTGTCGGCGCAGACGGCGGG |
| forE R4 | SPR oligo for forE with linker | CCCGCCGTCTGCGCCGACACCGGCGTCATCAACTTCATCGcctaccctacgtcctcctgc |
| forE F5 | SPR oligo for forE | GGCGCAGACGGCGGGCCTGGTCGCCGAACTCCTCGTACCG |
| forE R5 | SPR oligo for forE with linker | CGGTACGAGGAGTTCGGCGACCAGGCCCGCCGTCTGCGCCcctaccctacgtcctcctgc |
| forE F6 | SPR oligo for forE | GAACTCCTCGTACCGGACGGGGAACCGGTCGAGTTCGGGC |
| forE R6 | SPR oligo for forE with linker | GCCCGAACTCGACCGGTTCCCCGTCCGGTACGAGGAGTTCcctaccctacgtcctcctgc |
| forE F7 | SPR oligo for forE | CGGTCGAGTTCGGGCAGCCGCTGCTCGCCATCGAACCCGC |
| forE R7 | SPR oligo for forE with linker | GCGGGTTCGATGGCGAGCAGCGGCTGCCCGAACTCGACCGcctaccctacgtcctcctgc |
| forE F8 | SPR oligo for forE | CGCCATCGAACCCGCCTGAGCCTCAG |
| forE R8 | SPR oligo for forE with linker | CTGAGGCTCAGGCGGGTTCGATGGCGcctaccctacgtcctcctgc |

**Media used in this study:**

| **Media** | **Recipe (per litre)** | **Water** | **pH** |
| --- | --- | --- | --- |
| SFM | 20 g soy flour  20 g mannitol  20 g agar | Tap |  |
| MYM | 4 g maltose  4 g yeast extract  10 g malt extract  +/- 18 g agar | 50:50 Tap:deionised | 7.3 |
| LB | 10 g tryptone  5 g yeast extract  10 g NaCl (omitted when selecting with Hygromycin)  +/- 20 g agar | Deionised | 7.5 |
| 2YT | 16 g tryptone  10 g yeast extract  5 g NaCl | Deionised | 7.0 |
